# The evolution of pectate lyase-like genes across land plants, and their roles in haustorium formation in parasitic plant, *Triphysaria versicolor* (Orobanchaceae)

**DOI:** 10.1101/2024.12.04.626843

**Authors:** Huiting Zhang, Eric K. Wafula, Elizabeth A. Kelly, Itsuhiro Ko, Alan Yocca, Pradeepa C.G. Bandaranayake, Zhenzhen Yang, Daniel B. Steele, John Yoder, Loren A. Honaas, Claude W. dePamphilis

## Abstract

Parasitic plants in Orobanchaceae are noxious agricultural pests that severely impact crops worldwide. These plants acquire water and nutrients from their hosts through a specialized organ called the haustorium. A key step in haustorium development involves cell wall modification. In this study, we identified and analyzed the evolutionary relationships of *pectate lyase-like* (*PLL*) genes across parasitic plants and other non-parasitic land plant lineages. To support detailed examination of gene models and paralogous gene family members, we used published parasitic plant genomes, as well as a recently generated draft genome assembly and annotation of *Triphysaria versicolor*. One particular *PLL* gene, denoted as *PLL1* in parasitic Orobanchaceae, emerged as an important candidate gene for parasitism. Our previous comparative transcriptomic analyses showed that *PLL1* underwent neofunctionalization via an expression shift from floral tissues in non-parasitic relatives to haustoria in parasitic species. It belongs to the largest sub-clade of the PLL gene family, is highly upregulated in haustoria, and shows signatures of relaxed purifying selection and 15 individual sites with signatures of adaptive evolution. To explore its function in haustorium development, we manipulated *PLL1* expression in *T. versicolor*, a model parasitic species from Orobanchaceae, using direct transformation with the parasite and host-induced-gene-silencing (HIGS). For HIGS, we generated transgenic *Medicago* hosts expressing hairpin RNAs targeting the *PLL1* gene in *T. versicolor*. An average 60% reduction of *PLL1* transcript level was observed in both direct transformation and HIGS treatments, leading to an increased frequency of poorly adhered parasites with fewer xylem connections and a smaller proportion of mature haustoria. These findings demonstrate that *PLL1* plays a crucial role in haustorium development and suggest it as a promising target for managing parasitic weeds. Notably, the success of HIGS even before the establishment of a functional haustorium highlights the possibility of early intervention against parasitism.

## Introduction

Pectins, a family of complex polysaccharide polymers, are major components of primary cell walls and the middle lamellae of land plants. In higher plants, pectin functions in maintaining the mechanical and structural integrity of plant cell walls, contributes to intercellular adhesion (Ridley *et al*., 2001), and plays a role in plant defense (Starr & Moran, 1962; Eticha *et al*., 2005). Several classes of pectin degrading enzymes have been identified in bacteria, fungi, and plants. One of those is <u>p</u>ectate lyase (PL), an enzyme that cleaves the α- 1,4-linked galacturonic acid in homogalacturonan by beta-elimination. PLs are well documented to play key roles in invasive growth of plant pathogens, such as *Erwinia carotovora* and *Bacillus* spp., which acts as extracellular agents in soft rot disease (Collmer & Keen, 1986; Kotoujansky, 1987; Nakajima *et al*., 1999). In plants, *pectate lyase-like* genes (*PLLs*) play essential roles in various developmental and reproductive processes. During pollen tube growth, *PLL* functions by loosening pollen cell walls to initiate pollen tube emergence (Ohtsuki *et al*., 1995; Wu *et al*., 1996; Ischebeck *et al*., 2008; Jiang *et al*., 2014). Plant *PLLs* are also involved in the activation of plant defenses by releasing plant cell wall oligogalacturonides as defense elicitors in response to various plant hormones and stresses (Palusa *et al*., 2007). In addition, studies of *PLLs* in horticulture crops revealed their roles in the development of important crop traits (Uluisik & Seymour, 2020). *GhPEL76*, a *PLL* gene in cotton, regulates the organ elongation in Arabidopsis and fiber elongation in cotton (Sun *et al*., 2020). Gene silencing assays in tomato and peach shed light on the roles that *PLLs* play in fruit ripening and softening (Yang *et al*., 2017; Xu *et al*., 2022).

Haustoria, the feeding structures that are present in all parasitic plants, are evolutionarily novel organs in the plant kingdom. Several studies have shown that parasitic plants have recruited haustorial genes from ancestral gene functions in nonparasitic plant tissues to adapt to a parasitic lifestyle (Yang *et al*., 2015; Yoshida *et al*., 2016; Cui *et al*., 2016), including those involved in pectin modification and degradation processes (Yang *et al*., 2015; Kurotani *et al*., 2020; Jhu *et al*., 2021a,b; Bawin *et al*., 2023; Leso *et al*., 2023; Dixit & Upadhyay, 2023; Bradley *et al*., 2024). One such example is the *PLL* gene family (Yang *et al*., 2015). In a study conducted by Yang *et al*., comparison between *PLL* gene expression patterns in the parasites and their orthologs in nonparasitic plants suggested that ancestral genes in parasitic plants were commonly expressed in floral tissues and that the haustorial expression was gained through regulatory neofunctionalization after a genome duplication event (Yang *et al*., 2015). Haustorial specific *PLL* genes from three parasitic species in Orobanchaceae were proposed to be good candidates for functional assays (Yang *et al*., 2015) and could be potential gene silencing targets for parasitic weed management (Fernández-Aparicio *et al*., 2020). Dynamic regulations of cell wall modification enzymes, including PLLs, were also observed in another parasitic plant, *Cuscuta*, during haustorium development and parasite-host interaction (Jhu *et al*., 2021a; Bawin *et al*., 2023; Bradley *et al*., 2024). However, the role of PLL in parasitic plants has not been experimentally characterized.

Unlike functional characterization in model plant species where high-efficiency protocols have been developed and tested exhaustively, functional assays in parasitic plants remain a challenging endeavor. Early attempts to use Host Induced Gene Silencing (HIGS) gave ambiguous or negative results in *Striga* (de Framond et al 2007) suggesting that HIGS may not be worth pursuing further for functional studies or the development of parasite resistant host genotypes. However, one reason for these outcomes may have been the lack of extensive sequence data in *Striga* and a rigorous framework for selecting specific gene targets most likely to yield detectable phenotypic effects.

To address this, Zhang (2020) proposed a strategy for selecting high-confident candidate genes for functional analysis in non-model organisms using a combination of transcriptome data, gene family phylogeny, and molecular evolution tests. To qualify as a good candidate gene, several analyses are needed on the parasitic plant *PLL* genes, such as validation of gene models, analysis of gene expression patterns on a uni-gene level, and identification of paralogous genes with potential functional redundancy; all of which can be facilitated with a gene family analysis accompanied by detailed expression data. However, a comprehensive view of the evolutionary history of the PLL family is lacking. Sun and van Nocker (2010) used the Neighbor-Joining method to construct a *PLL* family phylogeny in Arabidopsis, which served as the baseline for more detailed phylogenetic studies of *PLLs* across land plants (McCarthy *et al*., 2014; Zheng *et al*., 2018) or for a specific lineage (Sun *et al*., 2018; Xu *et al*., 2022). In the aforementioned cases, however, only a limited number of species were selected and the phylogenetic relationships between some large portions of the gene family remained unresolved or lacked strong support. Additionally, inferred phylogenetic relationships from those studies conflict with each other. To gain a deeper understanding of the evolutionary history of the *PLL* gene family in plants, especially in parasitic plants, and to verify whether the *PLLs* identified by Yang *et al*. (2015) meet the requirements of good candidate genes for informed functional studies, a well resolved, robust phylogeny that includes parasitic plants is needed.

In this study, with a goal of investigating the evolution of *pectate lyase-like* genes in parasitic plants and examining the involvement of *PLLs* in haustorium development, we implemented a phylogenetic approach to study the *PLL* gene family across land plants and gene knockdown assays on a selected *PLL* gene in parasitic plants. A haustorially expressed *Triphysaria versicolor PLL* gene, *TrVeBC3_16311*.*1*, designated as *TrVePLL1*, was selected for functional assays in the model parasitic plant *T. versicolor*. Both transient transformation via hairy root transformation (Tomilov *et al*., 2006; Bandaranayake *et al*., 2010) and <u>h</u>ost induced <u>g</u>ene <u>s</u>ilencing (HIGS) assays (Bandaranayake & Yoder, 2013) were used to knockdown the expression of the target gene. Knockdown assays resulted in fewer mature haustoria with weakened haustorial connections, demonstrating the involvement of *TrVePLL1* in plant parasitism and indicating that the HIGS knockdown strategy could contribute to parasite control in agriculturally damaging parasites.

## Materials and methods

### Pectate-lyase like gene identification and initial phylogenetic inference

The following processes were accomplished using the PlantTribes2 (Wafula *et al*., 2022) tools in Galaxy (https://usegalaxy.org/) (The Galaxy Community, 2022), following the method from Zhang et al., (2022).

Genes (*Arabidopsis PLL* genes), genomes (*Cuscuta* spp., *Ipomoea* spp., *Lindenbergia philippensis*, and *Striga hermonthica*), and transcriptomes of interest (*T. versicolor* build 4, *S. hermonthica* build 4, *P. aegyptiaca* build 6, and *L. philippensis* build 2 obtained from the Parasitic Plant Genome Project II, PPGP II, http://ppgp.huck.psu.edu/) were processed to predict protein coding regions using *AssemblyPostProcessor* with TransDecoder (Haas, 2024). A list of genomes and transcriptomes and data sources can be found in Supplemental Table S1. The post-processed sequences were classified into the pre-computed orthologous gene family clusters (orthogroups) of the 26Gv2.0 orthogroup scaffold from PlantTribes2 with *GeneFamilyClassifier* using the dual BLASTp (Camacho *et al*., 2009) and HMMER hmmscan approach (Eddy, 2011). The classified sequences were then assigned with the corresponding orthogroups’ metadata including functional annotation and super orthogroup assignment. Annotations of orthogroups within the same super orthogroups under all classification stringency were investigated to identify missing *PLL* genes. A total of four orthogroups were identified as *PLLs* from the PlantTribes2 26Gv2.0 scaffold based on both super orthogroup and functional annotation (Supplemental Table S2).

Sequences classified into desired orthogroups were integrated into the scaffold gene models using the *GeneFamilyIntegrator* tool and then all merged into one fasta file for downstream process. Multiple sequence alignments were generated using *GeneFamilyAligner* using MAFFT’s L-INS-i algorithm (Katoh *et al*., 2002) and the corresponding coding sequences were forced onto the amino acid alignment. Sites present in less than 10% of the aligned DNA sequences, and sequences with less than 50% alignment coverage were removed with trimAL (Capella-Gutiérrez *et al*., 2009). Maximum likelihood (ML) phylogenetic trees were estimated from the trimmed DNA alignments using two methods. First, a RAxML tree (Stamatakis, 2014) was constructed using the PlantTribes2 tool *GeneFamilyPhylogenyBuilder* with default settings plus 1000 bootstrap replicates to estimate the reliability of the branches on the trees. Second, IQ-TREE2 (Minh *et al*., 2020) was used with default settings to infer phylogenetic trees. In addition, 2000 ultrafast bootstrapping and the -bnni flag were used for UFBoot tree optimization (Minh *et al*., 2013).

### Assembly and annotation of the Triphysaria versicolor genome

Genome assemblies have been shown to be useful resources to validate and improve gene models (Marx *et al*., 2016; Li *et al*., 2018; Pilkington *et al*., 2018; Zhang *et al*., 2022; Yocca *et al*., 2024). A draft assembly and annotation of the *Triphysaria versicolor* genome was used to facilitate gene model validation. This genome was first described by Daniel B. Steele as a part of a doctoral dissertation (Steele, 2021), where the material and methods were described in detail. A brief summary of the methods is provided below.

High-molecular-weight genomic DNA was isolated from frozen above-ground material of a single field collected *Triphysaria versicolor* individual using a modified CTAB method. The genomic DNA was size-selected for large fragment (>40 kb) enrichment on a BluePippin instrument (Sage Science, Beverly, MA, USA) and barcoded according to the instructions from 10X Genomics (Pleasanton, CA, USA). The resulting products were sheared to a peak size of 350 bp on an E220 Focused Ultrasonicator (Covaris, Woburn, MA, USA). The sequencing library was prepared using the 10X Genomics Chromium Genome Kit V2 and sequenced on an Illumina Hiseq X sequencer in PE150 mode (Illumina, San Diego, CA, USA).

The genomic sequencing generated 713.5 million paired-end reads. After barcode removal, the cleaned reads were input to supernova (v2.1.1) (Weisenfeld *et al*., 2017) and assembled using default parameters. The assembly was used for annotation with MAKER (v3.01.02-beta) (Holt & Yandell, 2011). Transcriptome sequences for *Triphysaria versicolor* generated using RNASeq data from the same dissertation work were provided as EST evidence. Protein sequences from *Mimulus guttatus* (*Erythranthe guttatus*) were acquired from Mimubase (http://mimubase.org/FTP/Genomes/Mguttatus_256_v2.0/, last accessed: May 27th 2025) and provided as protein homology evidence. A brief summary of genome assembly and annotation can be found in Supplemental Table 3.

### Manual curation of parasitic plant PLL genes and final phylogeny inference

Putative *Triphysaria versicolor PLL* transcripts from PPGP II were manually curated using webAUGUSTUS (Stanke *et al*., 2004) (https://bioinf.uni-greifswald.de/webaugustus/, last accessed May 7th 2024). Key input files for AUGUSTUS were prepared as follow.

#### The genome file

Each putative *PLL* transcript was aligned to the *T. versicolor* genome assembly using BLASTn to identify the corresponding genomic region. A genomic fragment spanning the target transcript plus 3 kb upstream and downstream was extracted with Geneious R9 and used as the genome file input.

#### The cDNA evidence file

Protein sequences predicted from each *T. versicolor* PLL transcript were searched against *PLL* orthogroup protein sequences, described above, using BLASTp. Sequences with the highest bitscore from closely related species (*Striga asiatica, Mimulus guttatus, Lindenbergia philippensis*, and *Solanum lycopersicum*) and Arabidopsis were selected as the *cDNA evidence file* input.

#### Other parameters

*Solanum lycopersicum* was selected as the model organism for AUGUSTUS’ *species* parameters. Genes from both strands were reported and analyzed.

Full-length genes were identified by aligning the original transcript, AUGUSTUS predicted gene model, and the reference sequences used as *cDNA evidence* for AUGUSTUS prediction. In cases where AUGUSTUS settings failed to predict full length genes, transcripts, predicted exons, and *Striga asiatica* gene were aligned to the *T. versicolor* genome assembly using the “*map to reference*” function in Geneious R9, and coding regions were manually determined with the *S. asiatica* gene as a reference. A final BLASTn search of the curated gene model against the genome assembly confirmed their genomic placement (Supplemental Table 4). Surprisingly, the *TrVeBC3_16311*.*1* gene reported by Yang et al., 2015 was missing in the *T. versicolor* transcriptome build 4 but was recovered by manual annotation using this method. The same approach was applied to refine *Cuscuta campestris* gene models, with *C. australis* and *Ipomea* orthologs as reference.

Using the curated gene sequences, multiple sequence alignment and phylogenetic analysis were repeated with the same methods described above. To provide a comparison with parasitic-inclusive datasets, PLL family trees were also constructed using only autotrophic plant sequences. In total, four phylogenies were generated: RAxML and IQ-TREE trees for autotropic plant only, and RAxML and IQ-TREE trees including both autotrophic and parasitic plants. These trees were used to infer monophyletic groups within the *PLL* phylogeny, which were named following the convention proposed by Sun and van Nocker (2010). Gene membership across clades were highly consistent among the four trees (Supplemental Table S5). Because clade topologies were identical and nearly the same genes were present in each clade, IQ-TREE phylogenies were selected for downstream analysis due to their higher bootstrap support.

### Domain prediction and amino acid frequency analysis

Domain structures of proteins in each clade were predicted using the command line version InterProScan (v5.59.91) (Jones *et al*., 2014) and searched against all the databases. To predict the presence of transmembrane domains and signal peptides, the protein sequences were submitted to the SignalP-6.0 (Teufel *et al*., 2022) server (https://dtu.biolib.com/SignalP-6, last accessed: April 2025) and TMHMM server v2.0 (Krogh *et al*., 2001) (https://services.healthtech.dtu.dk/services/TMHMM-2.0/, last accessed: April 2025). Default setting was used for TMHMM, whereas ‘Eukarya’ and ‘fast’ mode was selected for SignalP-6.0. The Pec_lyase_C domains predicted in each sequence by InterProScan were isolated and a MAFFT alignment was performed on the coding sequences comprising the focal domain. Sequences from the same clade identified from the gene family phylogenetic tree including all parasitic plants investigated in this research were extracted from the alignment and submitted to WebLOGO (https://weblogo.berkeley.edu/logo.cgi, last accessed: May 8th 2024; Crooks *et al*., 2004) to create the amino acid frequency plot.

### Selective constraint analysis

To infer adaptive or purifying selection, the EasyCodeML software (v1.41) (Gao *et al*., 2019) was used to run hypothesis testing using the CodeML program (Yang, 2007) from the PAML package. Because gene relationships were sometimes ambiguous in the full gene family tree, genes from individual clades and subclades were extracted and corresponding clade-specific trees were constructed following the same method described above. These clade trees are provided in Supplemental File 1. Within each clade, branches containing parasitic plant *PLL* genes and selected related non-parasitic asterid genes were identified for further analysis. Codon alignments were generated with MAFTT using gene sequences for the sequences from these selected branches and manually curated. Trees for each of the parasite-containing branches, hereafter termed “branch trees” were constructed using RAxML and the curated alignments.

The branch model in CodeML was applied to test for differences in selective pressure (dN/dS; ω) between genes designated as foreground and the rest of the tree, using nested models with preset parameters. Codon alignments and phylogenies corresponding to the selected branches were used as inputs, with parasite genes selected as foreground. A likelihood ratio test (LRT) was performed to compare the two ratio model (M2; parasites ω different from the background) against the one ratio model (M0; ω of both foreground and background are the same). For cases in which the relaxation of purifying selection was detected, a branch-site model was subsequently applied to identify individual codons under positive selection in the foreground lineages. This test used the preset nested model configuration and the Bayes Empirical Bayes (BEB) method (Yang *et al*., 2005) was deployed to estimate posterior probabilities (pp) of sites evolving under positive selection. Sites with pp < 0.80 were categorized as low probability, those with 0.80 <= pp < 0.95 as medium, and those with pp >= 0.95 as high.

For the branch containing *TrVePLL1*, the codon alignment was visualized and annotated in Geneious R9, enabling comparison of positively selected sites with the pectate lyase domain and predicted functional residues.

### PLL1 protein structure modeling

To assess whether amino acid substitutions in parasite-derived PLL1 proteins are associated with structural variation relative to non-parasitic species, protein models were generated for PLL1 homologs from *Triphysaria versicolor* (TrVePLL1), *Striga hermonthica* (StHeBC4_h_c11616_p_c9613), and *Lindenbergia philippensis* (LiPhi.10g047500.1.p). Full-length amino acid sequences were submitted to the AlphaFold (Abramson *et al*., 2024) Protein Structure Prediction server (https://alphafoldserver.com/; last accessed July 15, 2025) using default parameters.

Predicted 3D structures were first aligned pairwise using the align function in PyMOL (Schrödinger, LLC, 2015; https://github.com/schrodinger/pymol-open-source) to compare the overall structural similarity. Sites inferred to be under positive selection were then mapped onto the parasite-derived models. To better visualize structural differences near these sites within the conserved catalytic region, residues 130–370 were extracted from each model, and a second alignment was performed using the same procedure. As the full-length structures showed high overall similarity, with RMSD values of 0.300–0.319 Å across all pairwise comparisons, molecular docking was not pursued.

### Plant material

Seeds of *Triphysaria versicolor* were collected from an open pollinated population in a pastureland near Napa, California (GPS location: 38°13⍰33.2°°N, 122°16⍰11.7⍰⍰W; Wang *et al*., 2020). *Arabidopsis thaliana* Columbia-0 seeds were obtained from the Charles Anderson lab at the Pennsylvania State University (PSU). *Medicago truncatula* Jemalong A17 seeds were kindly provided by David Stout from USDA Agricultural Research Service, Pullman, WA and propagated in the Buckhout greenhouse at PSU.

### Seed germination

*T. versicolor* seeds were surface sterilized with 70% ethanol for 10 minutes with shaking and then washed twice with distilled water. Next, seeds were scarified with *Triphysaria* sterilization solution (TSS: 50% household bleach, 0.01% Triton X-100) for 30 minutes on a platform shaker at 95 rpm and then washed twice with distilled water. The treated *T. versicolor* seeds were transferred to petri dishes with plant growth medium (1/4 x Hoagland’s basal salt and nutrient mix, 7.5 g/L plant tissue culture grade agar, pH of 6.1), and sealed with Micropore surgical tapes (3M, Maplewood, MN, USA). Seeds were allowed to stratify for 3 days at 4°C in the dark before moving to a 16°C growth chamber with a 12-hour light regime under 30 μmoles/m2/sec light intensity.

*M. truncatula* seeds were scarified by soaking in 18M H2SO4 for 8 min. with gentle stirring every 2 minutes. After the seeds were thoroughly washed with reverse osmosis water five times, they were sterilized with TSS for 5 minutes with gentle shaking. The sterilized seeds were then washed with distilled water 10 times. Scarified and sterilized seeds were transferred to a new 50 ml tube with 20 ml distilled water and incubated on a shaker with gentle agitation at room temperature for 6 hours to overnight. Water was decanted after the incubation period and seeds were left in the tubes to grow in the dark for 12-24 hours at room temperature. The germinated seeds were transferred to petri dishes with the same growth medium used for *T. versicolor* and incubated in a growth room at 25°C with a 16-hour light regime under 100 μmoles/m2/sec light intensity.

*A. thaliana* seeds were sterilized using 30% bleach with 0.1% SDS for 20 min., and shaken with a vortex shaker every 10 min. The sterilized seeds were washed with distilled water five times and transferred to petri dish containing MS growth medium (1/2 x Murashige and Skoog basal salt mixture, 10g/L sucrose, and 8g/L plant tissue culture grade agar, pH of 5.7). The treated seeds were incubated in a 22°C growth chamber with a 16-hour light period.

### Parasite and host co-culture

*M. truncatula* and *A. thaliana* plants with robust root systems were transferred to co-culture medium (1/4 x Hoagland’s basal salt and nutrient mix, 7.5 g/L sucrose, 7.5 g/L plant tissue culture grade agar, pH of 6.1). Host roots were carefully arranged approximately along the radius of the 150 mm petri dish. *Triphysaria* root tips were placed 0.5-1 mm from the host roots. Co-culture plates were sealed with Micropore surgical tape (3M, Maplewood, MN, USA) and placed on growth racks with a 10-degree angle from level. The growth racks were placed in a growth room at 25°C with a 16-hour light regime under 100 μmoles/m2/sec light intensity until tissue collection. Note *Arabidopsis* was only used for the Triphenyl tetrazolium chloride staining assay (see Haustorium Staining Assays, below), while *M. truncatula* was used as the host for all other assays.

### Haustorium development assay and tissue collection

To track changes in *TrVePLL1* expression levels during haustorium development, wild type *Triphysaria versicolor* was co-cultured with *Medicago truncatula* for up to 10 days. Haustorium tissue was collected at 12, 24, 48, 72, 120, and 240 hours post inoculation (HPI). Pre-attachment haustoria (at 12 and 24 HPIs) were collected by making cuts adjacent to the swollen region of the parasite roots. The post-attachment haustoria (48-240 HPI) were harvested by making cuts in both the parasite and host roots within 1mm adjacent to the haustoria. Harvested tissue was immediately frozen in liquid nitrogen and stored at -80°C until processing.

### Plasmid construction

To construct the hairpin RNAi vector, a 503-nucleotide region near the 3’ end of the *TrVePLL1* open reading frame was amplified with primers containing four nucleotides, CACC, at the 5’ end of the forward primer. To prevent off-targeting in both the parasitic plant and the host plant, this fragment was searched against the *T. versicolor* transcriptome assembly build 4 and the host genome annotations (*i*.*e. Medicago truncatula* and *Arabidopsis thaliana*) using BLASTn. No perfect matches longer than 20 continuous nucleotides were found in any of the three targets. RNA isolation and cDNA synthesis was described below in the ‘RNA isolation and transcript level analyses’ section. The PCR product was recombined into the pENTR™/D-TOPO® entry vector using the pENTR™ Directional TOPO® Cloning kit (Invitrogen, Waltham, MA, USA) following the manufacturer’s protocol. The entry construct was confirmed by Sanger sequencing and sequence comparison with the reference. Next, a Gateway® LR recombination reaction was performed to transfer the target amplicon into the pHG8-YFP parent vector (Bandaranayke *et al*., 2010) facilitated by Gateway® LR Clonase® II enzyme mix (Invitrogen, Waltham, MA, USA). A double enzymatic digestion with restriction endonucleases XbaI and XhoI was performed to screen for positive hairpin vectors. The constructed vector is named *pHpTvPLL1* and was transformed into *Agrobacterium rhizogenes* strain MSU440 via electroporation. Primer sequences can be found in Supplemental Table S6.

### Hairy root transformation

*Triphysaria versicolor* transformation process was performed following Bandaranayake & Yoder, (2018) with modifications. *Agrobactrium rhizogenes* was grown on mannitol-glutamic acid: Luria-Bertani (MG/L) medium containing 400 uM acetosyringone (Sigma-Aldrich, St. Louis, MO, USA) overnight or until the bacteria covered the petri dish and formed a thick lawn. Ten to fifteen-day old *T. versicolor* seedlings were used for transformation. A clear cut was made at the joint between the hypocotyl and the root. The wounded hypocotyl end of the seedling was dipped into the *A. rhizogenes* culture several times. A petri dish containing Murashige and Skoog (MS) medium with 3% sucrose was treated with freshly prepared acetosyringone at the concentration of 400 uM. The *A. rhizogenes* infiltrated plantlets were transferred onto the aforementioned plates and incubated vertically in a 16°C growth chamber with a 12-hour light regime under 30 μmoles/m2/sec light intensity for 7 to 10 days. To initiate root growth, plantlets were transferred to new growth medium (1/4 x Hoagland’s basal salt and nutrient mix, 7.5g/L sucrose, 7.5 g/L plant tissue culture grade agar, pH of 6.1) with 300 uM Timentin and incubated for another 14 days. Transgenic calli and roots were identified by the visualization of YFP fluorescent signal using Zeiss SteREO Discovery.V12 with X-Cite® series 120 Q lamp under a YFP filter. Non-transgenic calli were discarded and non-transgenic roots were removed. Plants with positive signals were moved to a new growth medium and the tips of the transgenic roots were trimmed off to allow proliferation. Plates containing the transgenic plants were incubated vertically in a growth room at 25°C with a 16-hour light regime under 100 μmoles /m2/sec light intensity. Plants were screened and transferred to a new medium every 2 to 3 weeks until used for co-culture experiments. Because the transgenic *T. versicolor* roots have been through several rounds of root proliferation, an initial co-culture experiment was conducted with *T. versicolor* transformed with the empty vector and wild type *M. truncatula* to test their haustorium formation capability. Three to five plates were collected for each of the three biological replicates when using seedlings (control), and each plate contains 10 to 14 roots. Six to eight plates were collected for each replicate when using the transgenic line, and each plate contains three to eight roots with YFP signal. Haustorium formation rates were assayed five and seven days post co-culture. Statistical analysis is performed for this dataset and is described below.

Host plant, *i*.*e. Medicago truncatula*, transformation was performed following Boisson-Dernier *et al*., (2001) with modifications. *A. rhizogenes* were prepared as described above. *M. truncatula* seeds were germinated as described above and seedlings grown in the dark for 36-48 hours were used for transformation. To make a wound in the host plant seedling, the terminal 2-3 mm of the root tip was excised. The wounded plant was dipped in the *A. rhizogenes* culture and then transferred to growth medium (1/4 x Hoagland’s basal salt and nutrient mix, 7.5 g/L plant tissue culture grade agar, pH of 6.1) treated with freshly made acetosyringone at the concentration of 400 uM. The plates were placed vertically in the 16°C growth chamber for 5 days and transferred to fresh plates and incubated in the 25°C growth room. Transformed plants were subject to YFP screening 2 weeks post infiltration. Plants with no transgenic signal were discarded and roots with no YFP signal were removed. Plants were screened and transferred to a new medium every 2 to 3 weeks until used for co-culture experiments. To test for the presence of transformed DNA fragments, three types of *M. truncatula* roots were collected: 1) transformed roots with YFP signal, 2) transformed roots without YFP signal, and 3) wild-type roots. For each root type, three biological replicates were obtained. each containing 5-8 plants. The tissue was ground in liquid nitrogen and DNA isolated using the Macherey-Nagel (Düren, Germany) NucleoSpin® Plant II kit following manufacture’s protocol (section 5.1). Concentrations of the extracted DNA samples were assessed on the Qubit fluorometer (Thermo Fisher Scientific, Waltham, MA, USA) using dsDNA BR (broad range) Assay Kit. Same primers used to construct the RNAi construct were used to amplify a fragment of *TrVePLL1*. Gel electrophoresis was used to visualize the presence or absence of the amplicons.

### Haustorium categorization and tissue collection

To test the effect of *PLL* knockdown on haustorium development, co-culture experiments were performed on the following parasite-host pairs: 1) *T. versicolor* transformed with *pHpTvPLL1* and co-cultured with wild type host; 2) *T. versicolor* with *pHG8-YFP* (empty vector control) co-cultured with wild type host; 3) wild type *T. versicolor* co-cultured with a host transformed with *pHpTvPLL1*; and 4) wild type *T. versicolor* co-cultured with a host transformed with *pHG8-YFP*. Parasite-host co-cultures were monitored daily, and parasite root tips that initiated haustoria later than 24 hours post inoculation were excluded from analyses. In direct transformation experiments (parasite-host pairs 1 and 2), transgenic *T. versicolor* roots were grown with a wild type host for 7 days before harvesting. In host-induced-gene-silencing (HIGS) experiment (pairs 3 and 4), two-week-old *T. versicolor* seedlings were grown with transgenic host roots for 5 days before collection.

At the end of the co-culture period, the quality of the haustorial connection was tested by applying a gentle force: the host root was secured with one pair of tweezers and the parasitic root adjacent to the haustorium was pulled gently. Haustoria that detached easily from the host root were categorized as ‘failed’, while those remain attached were categorized as ‘mature’. To avoid potential observer bias, each each co-culture plate were assigned a random number, and the researcher recorded haustorium type without knowledge of the corresponding treatment.

Five or six biological replicates were created for each co-culture pair. In direct transformation experiments, 50-60 transgenic roots from 8-10 independent *T. versicolor* root cultures were used in each biological replicate. In HIGS experiments, 80-100 *T. versicolor* seedlings were inoculated onto 8-10 transgenic host roots in each biological replicates. Statistical tests were performed on the results and are described below. The co-culture and haustorium categorization assays were performed two more times at a smaller scale. Results from the three experiments are summarized in Supplemental Table S7.

To assay transcript levels of *TrVePLL1* in haustoria from co-culture experiments, failed and mature haustoria were collected. The failed haustoria were collected using the same methods for collecting pre-attachment haustoria. The mature haustoria were harvested using the same method for harvesting post-attachment haustoria. Three biological replicates were transferred immediately to liquid nitrogen for nucleic acid isolation. The rest were divided into two sets: one set for ink-vinegar staining and the other set for Toluidine Blue O staining (methods described below). Those samples were transferred immediately to the corresponding staining/fixation solutions.

### RNA isolation and transcript level analyses

RNA was isolated for cDNA synthesis in the plasmid construction process and for qPCR assays to estimate gene expression levels. Haustorial tissues collected for RNA isolation were ground in 350 ul RA1 buffer (Macherey-Nagel, Düren, Germany) mixed with 1% beta-mercaptoethanol (Qiagen, Hilden, Germany) using Kimble Chase glass tissue grinders (part# KT885450-0020). Total RNA was isolated from the macerated tissue using the Macherey-Nagel NucleoSpin® RNA Plant kit following the instructions from the user manual (section 5.1). For each sample, RNA concentration was assessed on the Qubit fluorometer (Thermo Fisher Scientific, Waltham, MA, USA) using the RNA HS Assay Kit, while RNA purity was assessed with a NanoDrop spectrophotometer (Thermo Fisher Scientific, Waltham, MA, USA).

cDNA was synthesized from RNA used for transcriptome sequencing in the PPGP project (Yang *et al*., 2015) or RNA collected for this project. 200 nanograms of RNA from each sample were converted to cDNA using the AzuraQuant™ cDNA Synthesis Kit (Azura Genomics, Raynham, MA, USA). The resulting cDNA was used for plasmid construction (described above) and transcript level estimation.

Primers for transcript level measurement were designed using Primer3 (v. 0.4.0, http://bioinfo.ut.ee/primer3-0.4.0/; Koressaar & Remm, 2007; Untergasser *et al*., 2012). To test primer specificity, melting curve analysis was performed on each PCR reaction and primer pairs yielding more than one peak in the melting curve were excluded.

The reverse transcription products were diluted 10-fold, and 1ul was used for each the qRT-PCR reaction with PerfeCTa SYBR Green FastMix (QuantaBio, Beverly, MA, USA). The reaction was performed with StepOnePlus Real-Time PCR System (Thermo Fisher Scientific, Waltham, MA, USA) with the following program: Step 1. 95°C for 5 min.; Step 2. 95°C for 0.5 min.; Step 3. 60°C for 0.5 min.; Repeat Step 2 and 3 for 40 cycles; Melt curve: 0.5 increments from 95°C to 25°C.

The average crossing point (Ct) values from three technical replicates were used to represent each biological replicate and three biological replicates were measured for each sample. The 2^(-ddCt) method (Livak & Schmittgen, 2001) was used to calculate the fold change in expression of target genes relative to the constitutively expressed gene *TrVePP2A* (Bandaranayake *et al*., 2010). Statistical tests were performed on the results and are described below.

### Haustorium staining assays

Three staining assays were deployed to examine morphological changes within the haustoria formed from the transgenic root and the control roots. Triphenyl tetrazolium chloride (TTC) staining was performed to observe metabolic activities in the haustoria (Stūrīte *et al*., 2005); Toluidine blue O (TBO) staining was performed to investigate changes in cell wall components in the haustoria (Pradhan Mitra & Loqué, 2014), while modified ink-vinegar staining assay was performed to visualize xylem bridges and other internal features of the haustorium. Samples for TBO and ink-vinegar staining assays were collected from the same experiment that categorized haustoria (see *Haustorium categorization and tissue collection*, above). A separate co-culture experiment was performed to collect tissue for the TCC staining assay. Three biological replicates were collected for each staining assay.

TTC staining was performed following Porter *et al*., (1947) with modifications. *M. truncatula* root accumulates red or green pigments in co-culture conditions, therefore, *Arabidopsis* was used in this experiment instead. 1% tetrazolium solution was prepared and stored at 4°C in the dark. Haustoria connected to the same host root were collected together and immediately transferred to a 50ml falcon tube containing 30 ml of 1% tetrazolium solution. The plant tissue was immersed in tetrazolium solution and then incubated at room temperature in the dark for 24 hours before screening. Zeiss SteREO Discovery.V12 was used to visualize the staining result and capture images.

TBO staining was performed on haustoria infiltrated with 4% paraformaldehyde. The Leica TP1020 - automatic tissue processor at PSU Core Microscopy Facility was used to dehydrate and infiltrate the tissue with paraffin (process described in Supplemental table S8). One to several haustoria were embedded in a paraffin mold with the host root oriented on the vertical axis using the Leica EG 115°C tissue embedding workstation. Sections with the thickness of 8*u*m were made with a rotary microtome (Thermo Fisher Scientific, Waltham, MA, USA) and de-waxed with a Leica autostainer XL with a program detailed in Supplemental table S8. TBO solution was added to regions on the slides where tissue was present and incubated at room temperature of 30 seconds. Slides were cleaned with distilled water. Stained slides were visualized, and images were captured with an Olympus BX61 microscope.

Ink-vinegar staining of the haustoria was performed following Vierheilig *et al*., (1998) with modifications. Host roots harboring one to several haustoria were collected in tissue embedding cassettes and placed in glass staining dishes with clearing solution (10% KOH solution). Samples were incubated at room temperature for 4 to 5 days and examined under dissecting scope daily to determine whether the clearing process was completed. Once the tissue is clear, cassettes were moved to a 1 L stainer and rinsed in cold running water for 5 min. The cassettes were then submerged in acidified water (5 ml white table vinegar in 95 ml water) for 5 minutes and rinsed briefly under cold tap water. The cleared and rinsed samples were placed in a preheated 5% ink solution (5 ml india ink in 95 ml water) in a water bath at 90°C for 30 min. Cassettes were removed to allow staining solution to drain, and then cassettes were rinsed under cold running water until water runs clear. Stained samples were moved to sterilized acidified water overnight to remove excessive ink staining. Samples were visualized and images were captured with Zeiss SteREO Discovery.V12.

### Statistical analyses

Due to limited sample sizes, both parametric and non-parametric statistical analyses were performed. For pairwise comparisons, a Mann-Whitney U test (non-parametric) and a Student’s t-test (parametric) were performed. For multiple group comparisons, a Kruskal– Wallis test (non-parametric) and a one-way ANOVA test (parametric) were performed. Tests for normality and equal variance were performed prior to the parametric test to validate the assumptions. Shapiro-Wilk test was used to assess normality, while F-test (for pairwise comparisons) or Bartlett’s test (for multiple groups) were used to evaluate equal variance. For the qPCR data, one-tailed p-values were used for directional changes (reduced gene expression in the knockdown lines), while two-tailed p-values were used for all the other analyses. Raw data and statistical results are available in Supplemental Table S9.

## Results

### Phylogenetic analysis of the pectate lyase-like gene family across land plants

To investigate the evolutionary relationships of *PLL* genes, Maximum Likelihood (ML) trees were constructed using two datasets. The first dataset included PLL genes from autotrophic land plants represented in the PlantTribes2 26Gv2.0 orthogroup scaffold, along with sequences from *Ipomoea nil* (Hoshino *et al*., 2016), *I. trifida (Wu et al*., 2018), and *Lindenbergia philippensis;* representing the nonparasitic sister group of the parasitic Convolvulaceae and Orobanchaceae, respectively. The second dataset expanded on the first by incorporating *PLL* genes from parasitic plants (species listed in Supplemental Figure S1 and data source listed in Supplemental Table S1).

Each dataset was used to construct a separate ML tree. The autotroph-only tree served as a reference to establish a baseline topology, as parasitic plant sequences can be highly divergent due to their unique lifestyle or less reliably annotated due to limited genomic resources and curation. In contrast, sequences from model autotrophic species tend to be more thoroughly curated, which helps reduce annotation-related artifacts that might otherwise distort branching patterns and decrease node support. The second tree, based on the combined dataset of autotrophs and parasitic species, was used to place parasitic *PLL* genes in a broader phylogenetic context and to compare them with homologs in their closest nonparasitic relatives.

Most of the five major clades identified in previous studies (Sun & van Nocker, 2010; McCarthy *et al*., 2014; Sun *et al*., 2018; Zheng *et al*., 2018) were strongly supported (bootstrap > 70) in both the autotroph-only and the combined autotroph-plus-parasite phylogenetic trees (Figure 1A, Supplemental Figure S2). Genes from all sampled angiosperm lineages were present in most clades. Clade 5, the earliest diverging monophyletic group (Figure 1A), contains genes from all land plants and forms the sister group to the remaining clades. Clades 4 and 3 form the second main monophyletic group, corresponding to seed plant-wide and eudicot-wide clades, respectively. These two clades are highly similar in sequence and form a well-supported monophyletic group along with two *PLL* genes from *Selaginella moellendorffii*.

**Figure 1.**
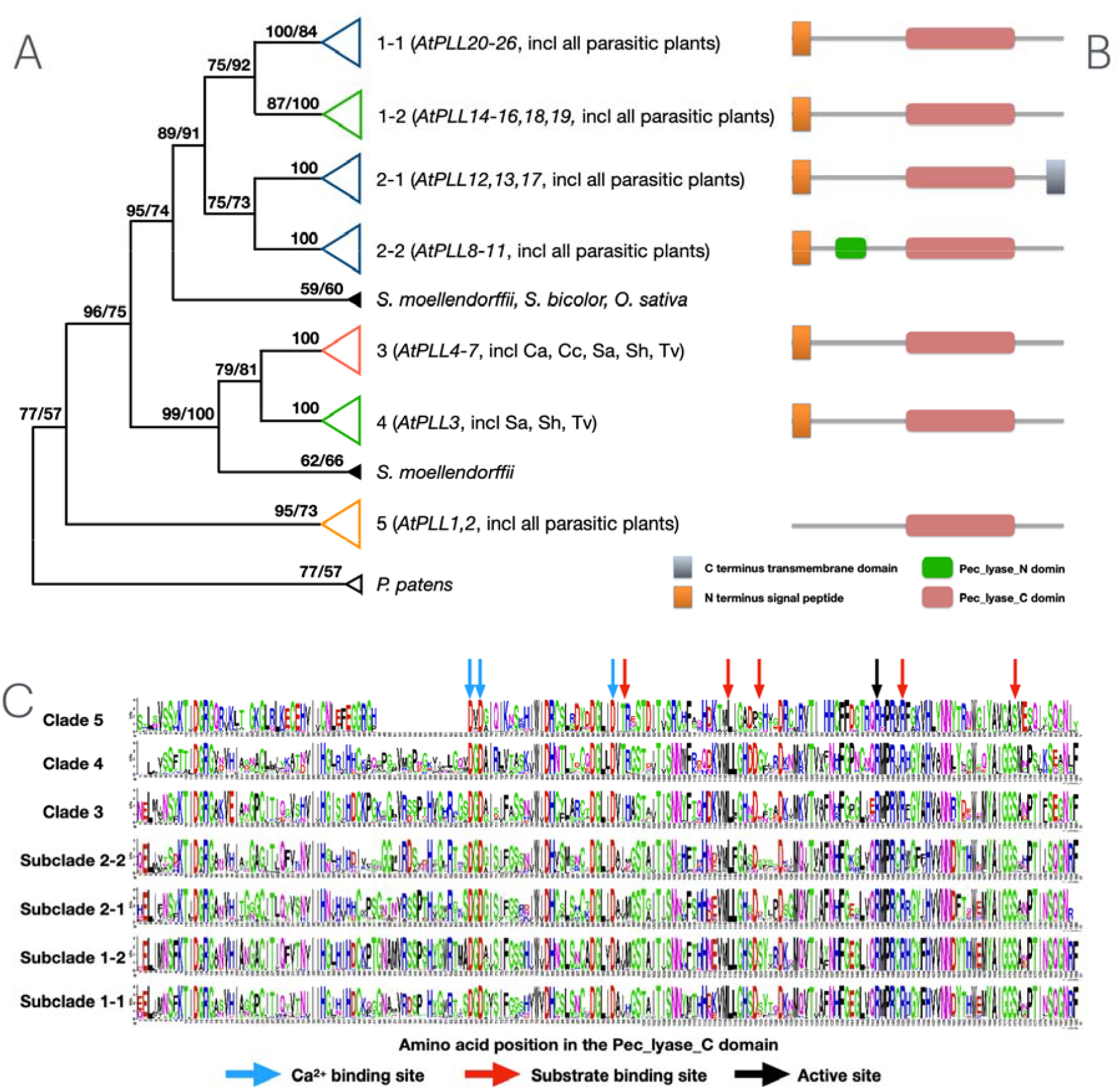
Phylogeny and domain structures of pectate lyase like (PLL) genes across land plants. (A). Simplified phylogenetic tree of the *PLL* gene family constructed with IQ-TREE. Monophyletic groups were represented with triangles and colored according to the range of species present in the clade. Blue: Angiosperm only; Green: Seed plant (Angiosperm + Pine); Pink: Eudicot only; Yellow: Land plant wide; Black: others. Numbers at nodes are bootstrap values. Arabidopsis genes housed in each clade are listed in the parentheses following the clade name along with abbreviations of parasites (Ca: *Cuscuta australis*; Cc: *Cuscuta campestris*; Pa: *Phelipanche aegyptiaca*; Sa: *Striga asiatica*; Sh: *Striga hermonthica*; Tv: *Triphysaria versicolor*). Numbers at the nodes are bootstrap values, where the number before ‘/’ is the value from tree constructed with only autotrophic plants and the number after is the value from tree constructed with both autotrophic and parasitic plants. The list of plants used for the analysis is available in Supplemental Figure S1 and data sources are listed in Supplemental Table S1. (B). Domain structure of proteins in each clade. (C). LOGO plot showing a comparison of amino acid frequency at equivalent positions of the Pec_lyase_C domain among the seven major clades and subclades. Arrows above the image indicate functional residues predicted based on published PLL protein structure from previous studies.

A notable expansion of Clade 4 *PLLs* was observed in most rosid species, but not in other major lineages, including the parasitic plants (Supplemental Figure S2). Furthermore, Clade 4 included *PLL* genes from *Striga* and *Triphysaria*, but none from *Cuscuta* or *Phelipanche*, in contrast to the other clades where all four parasitic lineages were represented. The grouping of Clade 3 and 4 has previously been observed in McCarthy *et al*., (2014), but not in the other publications.

Clades 1 and 2 comprise the third major monophyletic group. Each can be further divided into two subclades, 1-1 and 1-2, and 2-1 and 2-2. Subclades 1-1 and Clade 2 include *PLL* genes from all major angiosperm taxa, while Subclade 1-2 additionally contains genes from loblolly pine (*Pinus taeda*), indicating a broader seed plant-wide distribution. In contrast to previous phylogenies inferred using Neighbor-Joining methods (Sun & van Nocker, 2010; Sun *et al*., 2018; Zheng *et al*., 2018), where Subclade 2-2 appears as a sister to Clades 1 and 2, and Subclade 2-1 was classified within Clade 1 (Supplemental Figure S3), the present analysis provides strong support for the monophyly of Clade 2, a relationship originally suggested by McCarthy *et al*., (2014).

The haustorial *PLL* genes previously proposed for functional studies from the Yang *et al*., (2015) study were classified into Subclade 1-1 (Supplemental Figure S2 and Supplemental File 1), the largest clade in the *PLL* family. Additionally, these haustorial genes reside on a branch with a longer branch length than most other genes, indicating higher sequence divergence in the coding region. Expression analysis from the Parasitic Plant Genome Project II (PPGP II) transcriptome data (Supplemental Figure S4) revealed that among the four *Triphysaria PLL* genes in Subclade 1-1, *TrVePLL1* is the only one specifically expressed in haustorial stages, providing additional support for this gene as a good candidate for functional analysis.

### Conserved domain structure among PLL proteins

Domain search results (Figure 1B) showed that all sequences used in the phylogenetic inference contain a Pec_lyase_C domain, which is responsible for the primary catalytic function in pectate modification. In addition, proteins in Subclade 2-2 contain a unique Pec_lyase_N terminal domain. This domain likely originated in a common ancestor of angiosperms, as Subclade 2-2 includes representatives from all sampled angiosperms. Secretory signal peptides (SSP) were frequently detected at the N-terminus of proteins in Clades 1 through 4, but only a single Clade 5 gene, *Selmo_v1*.*0_412910*, harbors this domain (Supplemental File 2). This pattern suggests two possible scenarios: 1) this SSP was present in the common ancestor of all clades but lost in Clade 5, or 2) it was gained in the common ancestor of Clade 1-4 after divergence from Clade 5. Additionally, several proteins in Subclade 2-1 contain C-terminus transmembrane domains, suggesting a novel domain acquisition event by the ancestral protein of this clade.

An amino acid frequency plot (LOGO plot; Schneider & Stephens, 1990) was generated for the Pec_lyase_C domain from each clade to visualize sequence divergence (Figure 1C). Clade 5, the sister group to all the other *PLL* clades, exhibits an 18-amino acid deletion as well as additional sequence divergence relative to the others. Despite this divergence, key functional residues corresponding to putative Ca^2+^ binding sites, substrate binding sites, and the catalytic site, as predicted from previously characterized PL protein structures (Yoder & Jurnak, 1995; Scavetta *et al*., 1999; Vogel *et al*., 2002; Leng *et al*., 2017), were highly conserved in all clades, including parasitic plant sequences.

### Adaptive evolution in parasitic plant PLLs

Long branches in the phylogenetic tree, indicative of high sequence divergence, may reflect underlying biological processes such as adaptive evolution. To investigate potential shifts in selective pressure, a two-step analysis was performed on branches housing parasitic plant genes. First, a branch test was conducted with parasitic plant genes designated as the foreground and nonparasitic asterid genes as the background. Among the 21 branches analyzed, ten showed significant differences in selective constraint between foreground and background sequences. Of these, two branches exhibited relaxed purifying selection (Supplemental Table S10), including one that contains *TrVePLL1*, a gene proposed to be an ideal candidate for functional assays by Yang *et al*., (2015). In contrast, parasitic plant genes in the remaining eight branches showed stronger purifying selection, suggesting the proteins encoded by these genes are evolving more slowly than their orthologs in non-parasitic plants, possibly due to functional constraint or essentiality.

Second, branch-site tests were applied to the two branches with relaxed purifying selection to identify individual codons under positive selection. Both branches contain amino acid sites with signatures of positive selection in the foreground sequences (Supplemental Table S10). In the branch containing TrVePLL1, 15 amino acids were inferred to be under positive selection; of these, ten overlapped with the Pec_lyase_C domain, one overlapped a substrate-binding site, one is adjacent to a Calcium-binding site, and one is two residues from the active site (Figure 2A and Supplemental Figure S5). To evaluate whether these amino acid substitutions contributed to structural divergence, AlphaFold3 was used to predict the protein structures of PLL1 homologs from *T. versicolor, S. hermonthica*, and *L. philippensis*. Pairwise structural alignments revealed minimal differences, with root-mean-square deviation (RMSD) values ranging from 0.300 to 0.319 Å across all comparisons. Visual inspection of the aligned models further indicated that the positively selected sites did not result in substantial structural shifts (Figure 2B-D), suggesting that functional divergence in these parasite-derived PLL1 proteins may be mediated through more subtle effects, such as changes in surface chemistry or interaction interfaces, rather than large-scale conformational changes.

**Figure 2.**
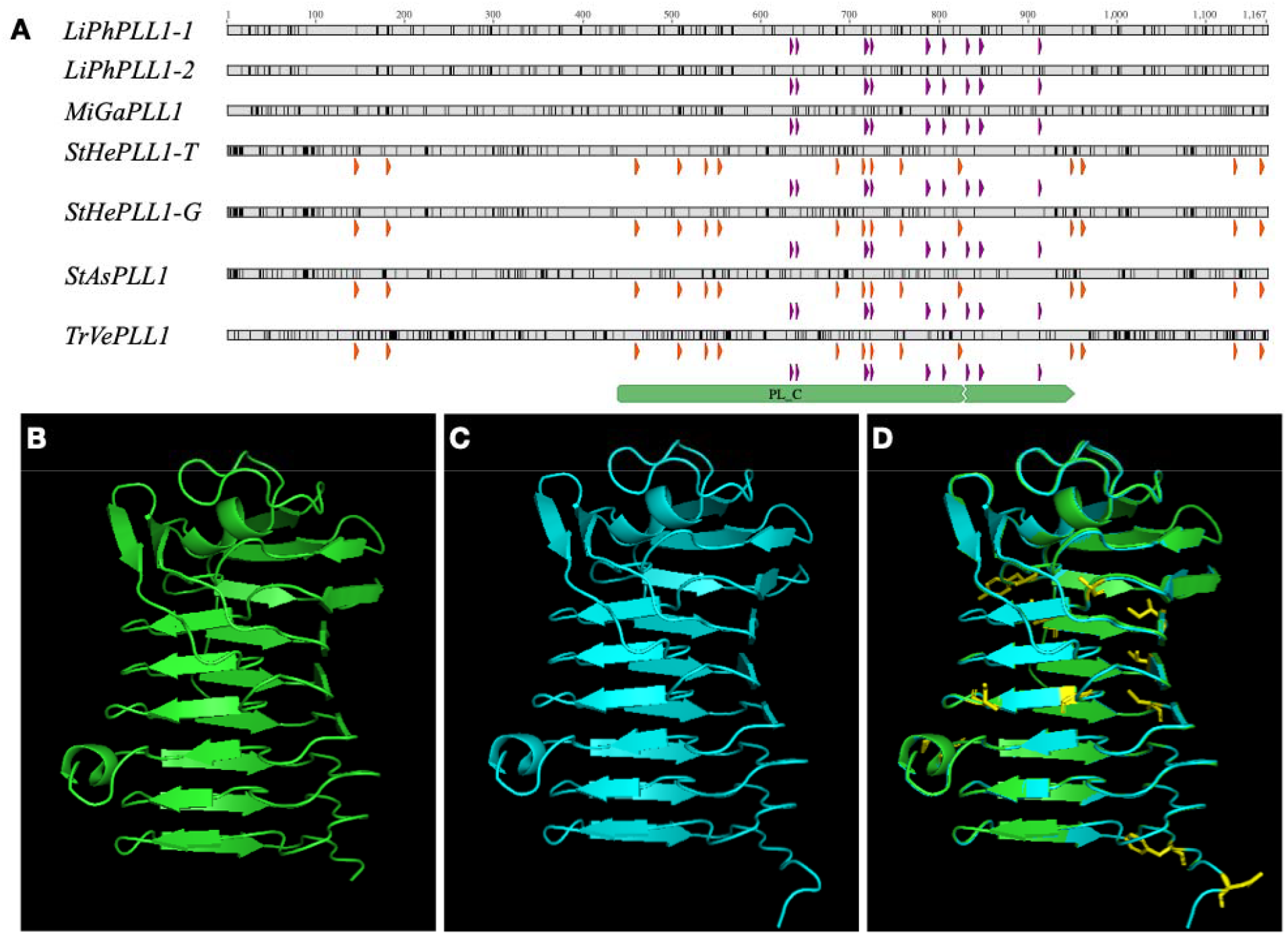
Structural and evolutionary analysis of PLL1 homologs. (A). Overview of the codon alignment from the PLL1 clade, showing the pectate lyase C domain (green bar), sites inferred to be under positive selection (orange blocks), and functional residues (purple blocks), including calcium-binding, substrate-binding, and the catalytic site. The full codon alignment with detailed residue-level annotations is provided in Supplemental Figure S5. (B) and (C) shows the AlphaFold-predicted structures of *Triphysaria versicolor* PLL1 (TrVePLL1) and *Lindenbergia philippensis* PLL1-1 (LiPhPLL1-1), respectively. (D) Structural alignment of TrVePLL1 (gray) and LiPhPLL1-1 (blue), with positively selected residues in TrVePLL1 shown as yellow sticks. The two structures are highly similar (RMSD⍰= ⍰0.30⍰Å).

### TrVePLL1 is upregulated during haustorium development

*TrVeBC3_16311*.*1*, a single *Triphysaria versicolor* gene, was chosen for functional assay and designated as *TrVePLL1*. This gene contains seven exons and six introns, encoding a 1338bp mRNA that translates into 446-amino acid protein (Figure 3A). Similar to proteins in Subclade 1-1, TrVePLL1 contains 3 functional motifs: an N-terminal signal peptide, a pectate lyase C domain, and a C-terminus low complexity motif (Figure 3A). Previous research demonstrated that *TrVePLL1* underwent neofunctionalization via an expression shift (Yang *et al*., 2015), and results from this study suggest additional evidence of adaptive protein evolution. These features make *TrVePLL1* a strong candidate for downstream characterization using RNAi-based molecular approaches.

**Figure 3.**
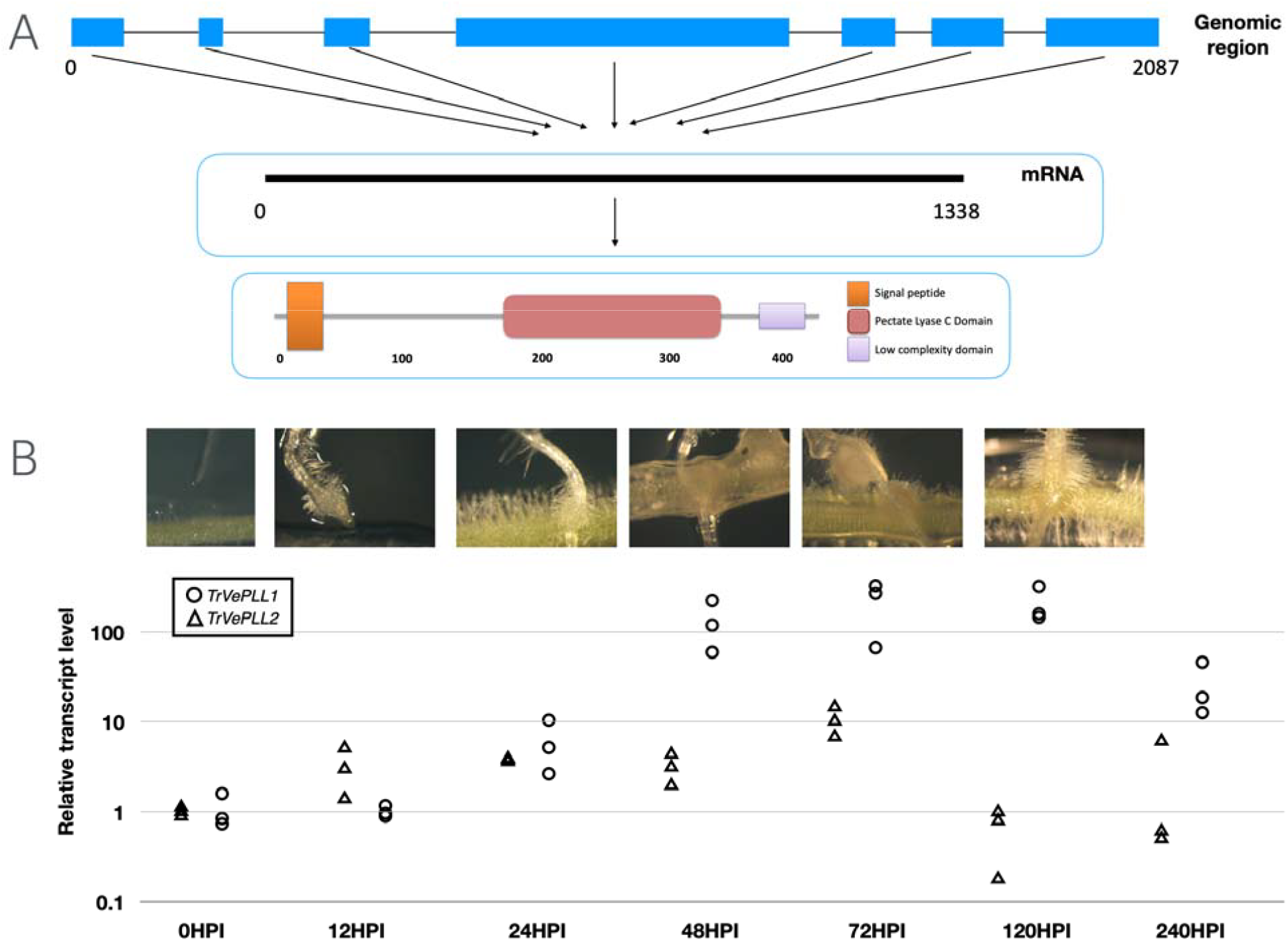
Gene model of TrVePLL1 and its expression levels during haustorium development in Triphysaria versicolor. (A). The seven exons (Blue box) transcribe into a 1341nt mRNA, which translates into the TrVePLL1 protein that contains three predicted functional domains: a signal peptide (Orange), a Pec_lyase_C domain (Pink), and a low complexity domain (Purple). (B). Haustorium development from 0 hour post inoculation till maturation (120 HPI, hours post inoculation) and relative transcript level of *TrVePLL1* and *TrVePLL2*. Total RNA of three biological replicates each consisting of haustoria from 6-8 plants were used for qRT-PCR. Note the log scale y axis.

Transcriptome data from PPGP (Yang *et al*., 2015) and PPGP II (accessible at http://ppgp.huck.psu.edu/) indicated that *TrVePLL1* is upregulated during early haustoria development (stage 3) (Supplemental Figure S4D). To validate this pattern and examine its expression profile at higher temporal resolution, transcript levels of *TrVePLL1* were measured from *T. versicolor* roots or haustorial tissues collected at 0, 12, 24, 48, 72 hours, as well as 5 and 10 days after incubation with *M. truncatula* roots (Figure 3B). Between 0 and 24 hours post inoculation (HPI), swelling occurred just behind the root tip of the parasite and the epidermal cells produced long haustorial hairs that adhered to the host root. Between 24 and 72 HPI, the haustorium apex extended toward the host and initiated penetration; during this phase, *TrVePLL1* expression increased sharply, reaching a plateau at ∼200-fold above baseline by 72 HPI. Elevated transcript levels persisted through 5 days post inoculation, coinciding with the formation of vascular connections, and declined by 10 days, when the haustorium had matured (Westwood *et al*., 2012).

Expression levels of *TrVeBC3_3448*.*1* (annotated as *TrVeBC4_p_c2216_g69* in PPGP II assembly, and designated as *TrVePLL2)*, was also measured in the same plant material as a control. Consistent with the transcriptome data, no significant expression changes were observed post inoculation. Two additional *TrVePLL* genes in Clade 1-1 were not tested due to very low transcript abundance in the PPGP II transcriptome datasets.

### Reduction of mature haustorium formation was achieved by knocking down TrVePLL1 via direct transformation and HIGS

To investigate whether *TrVePLL1* contributes to haustorium formation, RNA interference (RNAi) was used to knock down gene expression via two methods: 1) direct transient hairy root transformation of *Triphysaria versicolor*, and 2) host-induced-gene-silencing (HIGS), in which the silencing construct is transformed into the host plant.

A 503-base pair fragment of the *TrVePLL1* gene was cloned into the *pHG8-YFP* RNA silencing vector to generate the *pHpTvPLL1* hairpin construct. On average, 73% and 68% *T. versicolor* root transformed with the empty vector (*pHG8-YFP*) and *pHpTvPLL1*, respectively, showed YFP signals (Supplemental Figure S6), higher than previously reported efficiencies (Bandaranayake *et al*., 2010). Approximately 80% of the *M. truncatula* plants developed transgenic roots in both lines, consistent with previous publications (Bandaranayake & Yoder, 2013). The presence of the RNAi construct in *M. truncatula* roots was confirmed via PCR (Supplemental Figure S7). No morphological differences were observed between roots transformed with pHG8-YFP and pHpTvPLL1.

To generate sufficient root material roots for haustorium assays, transformed *T. versicolor* roots were maintained in tissue culture for up to three months with periodic root trimming to stimulate proliferation. These roots remained competent for haustorium formation, similar to non-transformed seedlings as shown in Supplemental Table 11.

In direct transformation experiments, both control and knockdown lines developed haustoria that appeared swollen, hairy, and attached to the host at seven days post inoculation. In the control lines, most haustoria are firmly connected to the host root, namely mature haustoria, while a small percentage (∼28%, Figure 4A) are loosely connected and detect readily from the host upon the application of a small mechanical force. Those loosely connected haustoria were deemed as ‘failed haustoria’. In contrast, the *TrVePLL1* knockdown lines exhibited a significantly higher proportion (∼45%) of failed haustoria. This phenotype was reproducible across three independent experiments (Supplemental Table S7).

**Figure 4.**
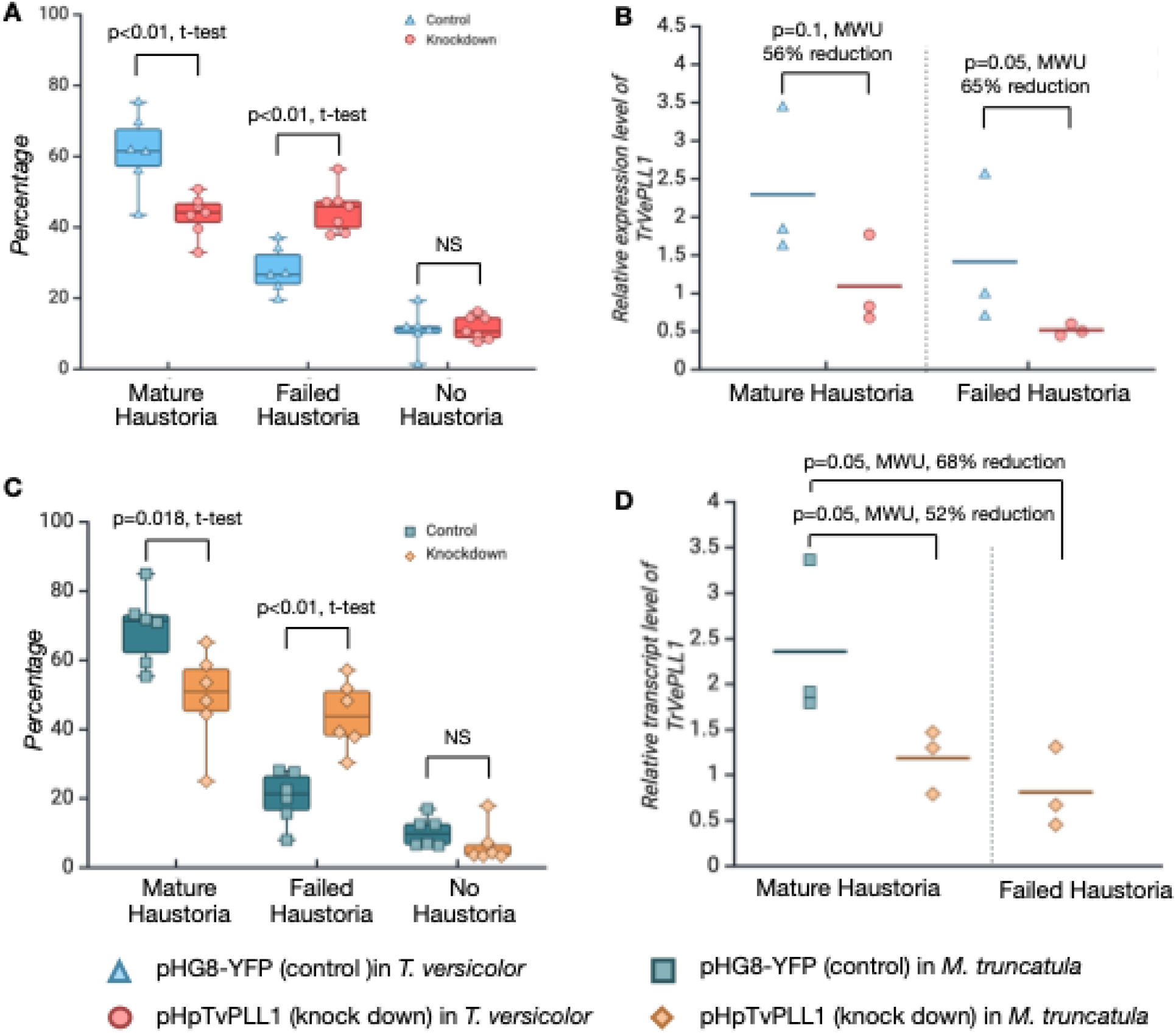
Haustorium formation rate and haustorium categorization in different co-culture experiments and quantification of *TrVePLL1* transcript level. *pHpTvPLL1* and the control vector *pHG8-YFP* were transformed to *Triphysaria* root (A) or *Medicago* root (C) using a hairy root transformation system. Six or seven biological replicates were used in (A) and (C), respectively. 50 to 60 transformed roots from 8-10 independent *Triphysaria* root cultures were analyzed in each biological replicate in (A). 80-100 wild type *Triphysaria* seedlings were co-cultured with 8-10 transformed *Medicago* roots in each replicate in (C). (B) and (D) show steady state transcript levels of *TrVePLL1* determined by qRT-PCR and normalized to the constitutively expressed gene *TrVePP2A*. 8 to10 haustoria were collected for each biological replicate and three biological replicates were generated for each treatment. Expression levels of *TrVePLL1* in *pHpTvPLL1* mature were set as 1. Horizontal lines among the data points represent the means. Statistical tests were performed and results are detailed in Supplemental Table S9. “t-test” stands for Student’s t-test, whereas “MWU” stands for Mann-Whitney U test. Note that no qRT-PCR was performed on failed haustorium from wild type *Triphasaria* co-cultured with *Medicago* roots transformed with *pHpTvPLL1* due to the small amount of failed haustoria collected from this co-culture experiment.

The same haustorium assay was repeated using a host-induced gene silencing (HIGS) system. In the control group, ∼69% of roots formed mature haustoria, and ∼20% formed failed haustoria (Figure 4C). When co-cultured with *M. truncatula* expressing *pHpTvPLL1*, only ∼49% of *T. versicolor* roots developed mature haustoria, while ∼44% produced failed haustoria. These results were highly consistent with those from the direct transformation assay.

To determine whether the increase in failed haustoria correlated with the expected reduction in *TrVePLL1* transcript level, a qRT-PCR assay was conducted. In the direct transformation system, *TrVePLL1* transcript levels were significantly reduced in both mature and failed haustoria from the knockdown lines compared to those from the control, with an average reduction of 60.5% in expression (Figure 4B). In the HIGS system, a similar trend was observed; however, failed haustoria from the control group were not sampled due to their low occurrence. In this system, both mature and failed haustoria from *T. versicolor* co-cultured with pHpTvPLL1-transformed hosts showed significantly reduced *TrVePLL1* transcript levels relative to the control, averaging a 60% reduction in expression (Figure 4D).

### Staining of haustoria reveals morphological similarities and differences between control and knock down lines

It is difficult to visually distinguish failed from the mature haustoria, or to differentiate a haustorium formed in the knock down lines from those in the control lines. To examine potential differences in internal haustorium structure, three histological staining assays were performed on haustoria collected from transgenic *Triphysaria versicolor* co-cultured with wild type hosts.

The ink-vinegar staining method revealed the presence of one or more xylem connections (Figure 5A, B) in most haustoria. Over 70% of mature haustoria developed single or multiple xylem bridges, while only about 50% of failed haustoria developed xylem connections (Figure 5C, and Table 2). Statistical analysis was conducted to assess differences in xylem bridge formation among four haustorium groups: mature and failed haustoria from both control and *TrVePLL1* knockdown lines. For the first comparison, which evaluated the percentage of haustoria that formed at least one xylem bridge, the failed haustoria from the control group were excluded due to low sample size. A Kruskal–Wallis test yielded a p-value of 0.063, indicating a marginal difference among the remaining groups. To further examine this trend, a one-way ANOVA was performed and revealed a significant difference (p = 0.013). Post hoc Tukey’s HSD test showed that the frequency of xylem bridge formation in failed haustoria from the knockdown line was significantly lower than in both mature haustoria from the same line and mature haustoria from the control line (Supplemental Table S9D). In the second comparison, which evaluated the average number of xylem bridges per haustorium, all four groups were included. No significant differences were observed among them (Table 1; Supplemental Table S9E).

**Table 1.** Xylem connection observations in transgenic haustoria. Statistical analysis results can be found in Supplemental Table 9D and E.

| Type of haustoria | % roots with xylem bridge<sup>a</sup> | Average number of xylem bridges in a haustorium<sup>b</sup> |
| --- | --- | --- |
| pHG8-YFP mature | 67 ± 6 (n= 62) | 1.32 ± 0.85 (n=41) |
| pHG8-YFP failed | 53 (n= 17) | 1.30 ± 0.67 (n=10) |
| pHpTvPL1 mature | 72 ± 8 (n= 46) | 1.28 ± 0.79 (n=39) |
| pHpTvPL1 failed | 50 ± 4 (n= 40) | 1.15 ± 0.50 (n=20) |

a Value are means ± standard deviation from three biological replicates (except for *pHG8-YFP* failed). Three to five plates were collected for each replicate. n = the total number of roots in all replicates. b only haustorium formed xylem bridge were included in the calculation

**Figure 5.**
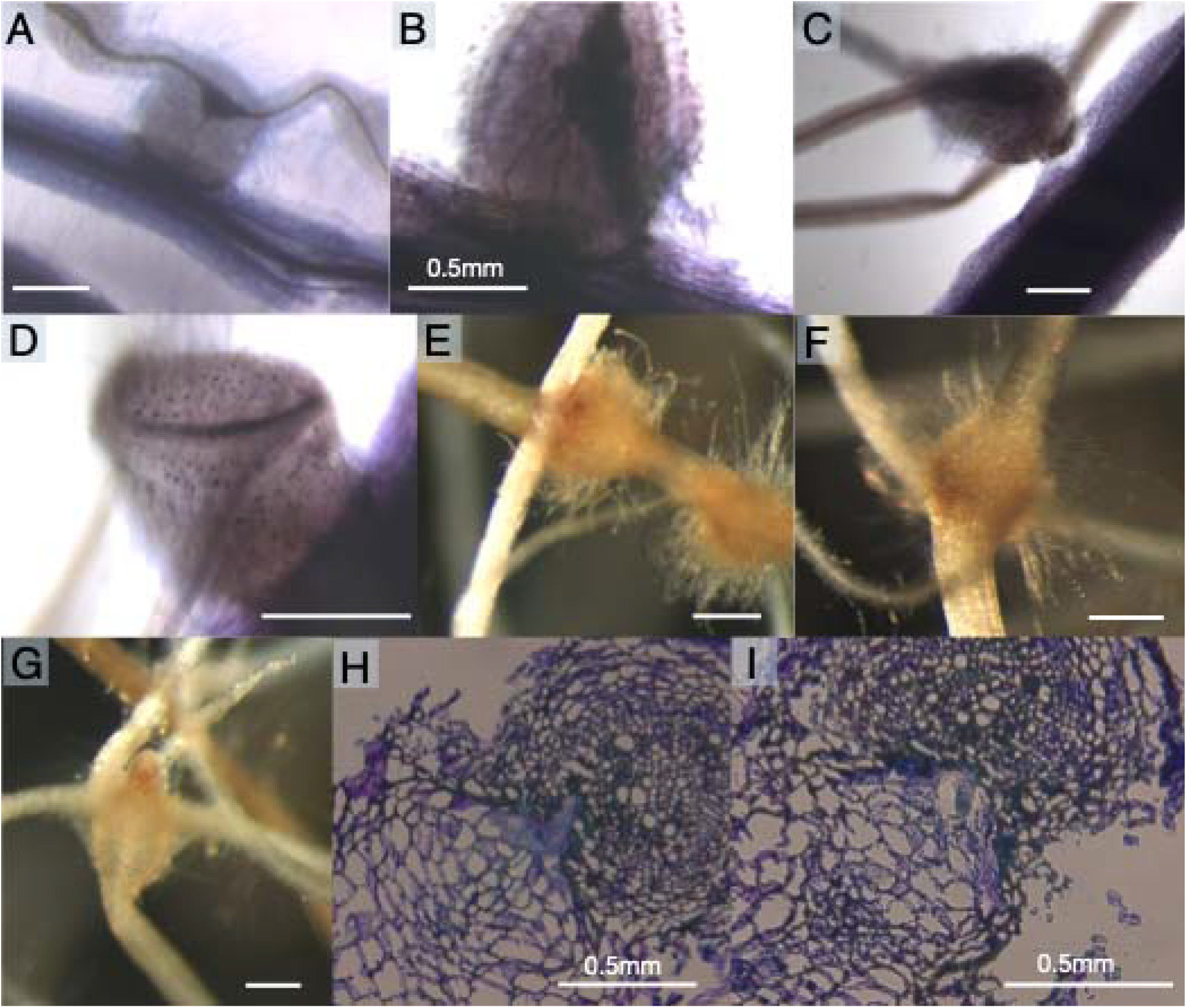
Morphological differences and similarities among haustoria formed by transgenic *Triphysaria* roots. (A)-(D): Ink and vinegar stained haustoria at 7 days post inoculation (DPI), showing difference in xylem connection. (E)-(G): Tetrazolium stained haustoria at 7 DPI. (E) and (F) showing a darker color in the parasite root, especially near the haustorial apex. (G) showed color accumulation at the parasite root tip. (H) and (I): 7 DPI haustorium sections stained with Toluidine blue O (TBO). (A), (C), (D), (F), (I) were collected from plants transformed with the hairpin construct, pHpTvPL1, whereas (B), (E), (G) and (H) were collected from plants transformed with the control vector, pHG8-YFP.

In the Triphenyl tetrazolium chloride (TTC) staining assay, haustoria were stained to a deep red, especially at the host-parasite interface (Figure 5E, F), consistent with high metabolic activity. In some instances, intense staining was also observed at root tips (Figure 5G). No clear differences were observed between failed and mature haustoria, or between the knockdown and the control lines in terms of TTC staining patterns.

Toluidine blue O (TBO) staining, distinguishes cell wall components based on acidity and chemical composition (Pradhan Mitra & Loqué, 2014), was used to examine histological features in sectioned haustoria. However, haustorium sections processed from the *TrVePLL1* knock down line and the control line were not visibly different (Figure 5H, I). It’s worth mentioning that only mature haustoria were used for sectioning in the TBO study, as failed haustoria frequently detached from the host root during the dehydration and embedding process.

## Discussion

### A robust phylogenetic analysis reveals the evolutionary history of the PLL genes

Compared to the published phylogenetic estimation of the *PLL* gene family (Palusa *et al*., 2007; Sun and Nocker, 2010; McCarthy *et al*., 2014; Zheng *et al*., 2018; Sun *et al*., 2018), the results from this research are both more thorough and robust. The Bayesian tree constructed by Palusa *et al*., (2007) divided 62 *PLL* proteins from *Arabidopsis thaliana, Populus trichocarpa*, and *Oryza sativa* into 3 distinct groups. With rice as the only monocot representative in that study, group II and a few other subgroups were annotated as dicot-specific. However, with increased sampling from diverse monocot species, the phylogeny presented in this research identified monocot genes in these groups. The only truly dicot specific clade is Clade 3, which was identified as the *PLL4* subgroup in group I from the Palusa et al. results. McCarthy *et al*. (2024) expanded the species list from three to ten and built a high-quality phylogeny to infer the ancestry of the *PLL* gene family. The main limitations in the McCarthy *et al*. phylogeny were the lack of resolution between some major clades and the absence of asterid representatives due to the then available genomes. By sampling a wider range of plant species, and including genes from eight asterid genomes and transcriptomes, all the clades in this study were resolved into monophyletic groups with strong support, with little ambiguity remaining in terms of the relationship between major clades. There have been other attempts to resolve the phylogeny of PLLs across land plants with additional plant genomes including asterids (Sun *et al*., 2018; Zheng *et al*., 2018). However, both publications failed to provide supporting evidence for the clade-to-clade relationship as we do in this study. By resolving all major clades with strong support and expanded taxon sampling, this phylogeny provides a critical framework for interpreting domain evolution, gene expression, and lineage-specific adaptation.

### How does TrVePLL1 facilitate parasitism? Two hypotheses

In the process of pollen tube growth, pectin deposition and the action of several pectin processing enzymes have been proposed to play essential roles. As a fast-growing tissue that penetrates through other plant tissues, the apical cell walls of the pollen tube must remain flexible enough to allow incorporate new cell wall material, while also maintain sufficient strength to withstand turgor pressure (Gaffe *et al*., 1994). As major cell wall components, pectic molecules have been reported to be actively delivered to the cell surface via vesicles (Dashek & Rosen, 1966; Rosen & Gawlik, 1966; Heslop-Harrison & Heslop-Harrison, 1982; Ischebeck *et al*., 2008). These pectic molecules may be newly synthesized and secreted from the Golgi apparatus, or recycled from older cell walls that have been degraded by pectic enzymes (Wu *et al*., 1996). In parallel, pectate lyase-like proteins have also been proposed to loosen the cell wall of the stylar cells, facilitating pollen tube penetration (Wu *et al*., 1996).

Haustorium penetration, which is morphologically similar to pollen tube growth, may have adopted analogous molecular strategies. In fact, some *PLLs* in the parasitic Orobanchaceae are derived from ancestral genes formerly expressed in floral tissues of their non-parasitic relatives (Yang *et al*., 2015). In addition, Losner-Goshen *et al*., (1998) used immunolabeling with pectin-specific antibodies and observed the disappearance of pectins from the cell walls of a sunflower root at the parasite-host interface, while the host cells themselves stayed intact. More recently, Auriac *et al*., (2024) provided additional support for the idea that parasitic plants penetrate host tissues intracellularly, rather than disrupting host cells directly.

Taken together, we propose two possible mechanisms by which *TrVePLL1* may function during haustorium development. *Hypothesis 1*: *TrVePLL1* facilitates the mobilization of parasite-derived pectin molecules by degrading demethylated pectin from older parasite cell walls, generating monomers that can be recycled as building material for new walls to grow at the apical tip of the haustorium. *Hypothesis 2*: *TrVePLL1* promotes haustorial penetration in the host tissue by targeting and degrading pectin within the host cell wall or middle lamella. To evaluate these hypotheses, further experiments, such as high-resolution tracking of TrVePLL1 protein localization or ultrastructural analysis of pectin content at the host–parasite interface, will be required.

### Working towards durable resistance in a changing environment

Despite the high heterozygosity level of gene sequences in the obligate outcrossing species *Triphysaria versicolor*, RNAi-based knockdown successfully reduced transcript level approximately 60% in both direct transformation and HIGS assay, with corresponding increase in failed haustoria. This success contrasts with earlier HIGS attempts in *Striga* (de Framond *et al*., 2007), where RNAi constructs were based directly on Arabidopsis “housekeeping” or “killer” genes essential for developmental or metabolism, rather than on homologous *Striga* sequences. Moreover, those efforts rarely assessed whether target genes were strongly expressed in haustoria, nor did they confirm reduced transcript levels in HIGS plants; these factors may have contributed to their weak or inconsistent phenotypic effects. Later HIGS studies (Runo *et al*., 2011) that used parasite-derived sequences, such as in *Triphysaria* (Bandaranayake & Yoder, 2013), *Phelipanche* (Aly *et al*., 2009), and *Cuscuta* (Jhu *et al*., 2021b), achieved effective gene silencing, demonstrated reduced transcript levels in HIGS lines, and highlighted the importance of using parasite sequences for successful knockdown. In this study, *TrVePLL1* was prioritized based on its strong and specific expression during haustorial development, a pattern conserved among orthologs in multiple parasitic lineages, suggesting an essential role in parasitism and a higher likelihood of effective silencing through RNAi. Moreover, evolutionary analyses identified *TrVePLL1* and its orthologs as lineage-specific and rapidly evolving, reducing the chance of off-target effects in non-parasitic plants. Together, these results highlight the potential of biologically informed, parasite-specific targeting strategies to improve the reliability and specificity of HIGS-based control approaches for weedy parasitic plants such as *Striga*.

Building on these findings, functional characterization of *TrVePLL1* orthologs in *Striga hermonthica* and *S. asiatica* (designated *StHePLL1* and *StAsPLL1*) should be prioritized if genetic manipulation becomes feasible, particularly as *StHePLL1* exhibits strong haustorial expression, high sequence divergence, and domain conservation, features that mark it as a strong candidate for parasitic weed control. Knockdown effectiveness could potentially be improved by incorporating insights from a growing understanding of genetic variation in target gene sequences, informed by population genetics surveys of *Striga* (Bellis *et al*., 2020). Such information could help design RNAi constructs that account for allelic diversity and maximize silencing efficiency across parasite populations.

Nevertheless, targeting a single gene may not confer durable resistance. In *T. versicolor*, 27 *PLL* genes were identified, including several that are expressed in the haustorium (Supplemental Figure S8). Although *TrVePLL1* is the only haustorially expressed member of Subclade 1-1 and exhibits features that distinguish it from other family members, partial functional redundancy may have contributed to the moderate knockdown phenotype, in which about 50% of haustoria still formed stable host connections. To address this limitation, future HIGS efforts may benefit from a multi-target “stacked” silencing strategy, as previously proposed by Gressel *et al*., (2017), in which RNAi constructs targeting multiple parasitism-related genes are combined to achieve broader and more durable suppression.

## Conclusions

Our results provide novel insights into the evolutionary history of the *pectate lyase-like* (*PLL*) genes in land plants and identified *TrVePLL1* as a key gene involved in haustorium development in parasitic Orobanchaceae. A phylogenetics tree constructed from *PLL* sequences across 34 plant species representing major groups of land plants, including six parasitic plants, revealed well-supported relationships among five major clades. *TrVePLL1*, a gene previously proposed as a candidate for parasitism-related function (Yang *et al*., 2015), was the only haustorial-specific gene in Subclade 1-1. Molecular evolution analyses further showed that the branch containing *TrVePLL1* exhibits signatures of relaxed purifying selection in parasitic lineages with 15 sites under positive selection, supporting its potential functional divergence. Together, these findings highlight *TrVePLL1* as a strong candidate gene for functional assays. Knockdown of *TrVePLL1* expression through transforming *T. versicolor* root with an antisense hairpin construct, or via host-induced gene silencing (HIGS), reduced the formation of mature haustoria. These findings provide some of the first experimental evidence implicating a cell wall-modifying enzyme in haustorium development (Jhu *et al*., 2021b; Leso *et al*., 2023), reinforcing the relevance of parasite-specific gene targeting strategies. With the continued advancements in genome and transcriptomic resources and improvements in computational tools, scientists are now well-equipped to identify additional candidate genes potentially involved in plant parasitism. However, functional validation remains a key bottleneck. Continued research, such as the study presented here, is essential for elucidating the molecular mechanism of parasitism and advancing new strategies for sustainable parasitic weed control

## Acknowledgements

The authors would like to thank Paula Ralph for technical support, and Prof. Charles Anderson for valuable methodological suggestions and access to microscopy equipment. This research is supported by NSF DBI-1238057 awarded to CWD and JY, with the additional support provided by the Plant Biology and Biology Department graduate programs at Penn State (for HZ, YZ, and LAH), the US Department of Agriculture, Agricultural Research Service (for HZ), Dr. Michael D. Frohlich (for HZ), and the Dorothy and Lloyd Huck Endowment (to CWD). Support for DBS was provided by UC Davis Plant Sciences Department GSR and the Jastro Shields Graduate Research Award.

## Completing interest

The author(s) declare no conflict of interest.

## Author contributions

CWD and HZ conceived the study. CWD and JY acquired funding. HZ, EKW, ZY, and LAH performed gene family analysis. HZ performed the protein structure analysis and selective constraint analysis. HZ constructed the plasmids, carried out transformation assays, haustorium assays, qPCR tests, and statistical analyses with assistance from EAK, IK, AY, and PCGB. DBS and JY assembled and annotated the draft *Triphysaria versicolor* genome. All authors contributed to the writing and editing of the manuscript.

## Data Availability

The *Triphysaria versicolor* genome assembly and annotation are available at FigShare (DOI: 10.6084/m9.figshare.29586506). All other data is available as supplemental materials.

## Supplemental Figure captions

**Supplementary Figure S1. Species tree of sampled plant taxa in the phylogenetic study**. Star (*) indicates parasitic plant species used in the research. Green box presented an alternative species relationship in the Orobanchaceae family, the common ancestor of which was marked with a yellow star. Different colors represent different major land plant groups. Green: asterid; Red: Rosid; Purple: Basal eudicot; Yellow: Monocots; Blue: Basal angiosperm and gymnosperm; Black: Lycophyte and Moss.

**Supplementary Figure S2. Phylogenetic trees of the PLL gene family constructed with only autotrophic plant PLL genes using IQ-TREE (A) or RAxML (B) and with PLL genes from both autotrophic and parasitic plants using IQ-TREE (C) or RAxML (D)**. Different colors represent different major land plant groups. Green: asterid; Red: Rosid; Purple: Basal eudicot; Yellow: Monocots; Blue: Basal angiosperm and gymnosperm; Black: Lycophyte and Moss.

**Supplementary Figure S3. Comparison of PLL gene family trees and gene family analysis methods from various publications and this work**. (A) Phylogeny from Sun and van Nocker, 2010; (B) Simplified phylogeny from McCarthy et al., 2014; (C) Simplified phylogeny from Sun et al., 2018; (D) Simplified phylogeny from Zheng et al., 2018; and (E) Simplified phylogeny from this work. (F) Summary and comparison of gene family analysis methods from the selected publications. Phylogenies in (B) to (E) are simplified to only show the overall structure with *Arabidopsis* genes for comparison purposes. Bootstrap (bs) values at the nodes represent the bs value of the largest monophyletic group housing the *Arabidopsis* genes shown at the tip. (C) and (D) lack bs values from the original publications. Colored backgrounds indicate clades identified by the Sun and van Nocker, 2010 work, whereas clades on the right side of each phylogeny shows the corresponding clades identified by this study.

**Supplementary Figure S4. Normalized transcript level (Transcript per Million, TPM) of *T. versicolor* genes from *PLL* Subclass 1-1**. Triplicated PPGP II transcript data was used for (A) to (C), and PPGP I transcript data was used for (D). X axis showing developmental stages (Stage abbreviated as S in the x axis legend). Blue boxes highlighting haustorial stages. *TrVePLL1* is the only *PLL* with principally haustorial expression.

**Supplementary Figure S5. Codon alignment for the branch containing *TrVePLL1***. Each gene is represented by two rows: the upper row shows the codon alignment, and the lower row shows the corresponding translated amino acid sequence. Nucleotides or amino acids that differ from the consensus are highlighted using the Clustal color scheme. The pectate lyase C domain was annotated below the alignment with a green bar. Sites inferred to be under positive selection are marked with orange blocks, where L in the block indicates low posterior probability, M indicates medium, and H indicates high. Functionally important sites were marked with purple blocks, including Calcium-binding sites (Ca), substrate-binding sites (S), and the active site (A).

**Supplementary Figure S6. *Triphysaria versicolor* (A-F) and *Medicago truncatula* (G-J) transformed with pHG8-YFP**. (A), (C), (D), (E), (G), and (I) are epifluorescence photos showing YFP expression. (B), (F), (H), and (J) are bright field photos. (A) and (B) showing a callus expressing YFP signal 2 weeks post *Agrobacterium* inoculation. (C) showing a single root developed from the transgenic callus 3 weeks post *Agrobacterium* inoculation. (D) showing transgenic root proliferation after removing root tip. (E) and (F) showing transgenic *Triphysaria* root forming haustorium on *Medicago* root 5 days post co-culture. (G) and (F) showing 6-week-old transgenic *Medicago* root without parasitization by *Triphysaria*. (I) and (J) are *Triphysaria* roots making haustorium on transgenic *Medicago* roots. C: callus; T: *Triphysaria versicolor* root; M: *Medicago truncatula* root; H: haustorium. Bar: 1mm, unless otherwise indicated.

**Supplementary Figure S7. Polymerase chain reaction of *TrVePLL1* using DNA from *Medicago truncatula* roots**. TY: transgenic root showing YFP signal; TnY: transgenic root not showing YFP signal; P: positive control, in which the *pHpTvPLL1* construct was used as template; NTC: no template control; nT: non transgenic root. Ladder: from the top to bottom band size: 650bp, 500bp, 400bp, 300bp, 200bp, 100bp.

**Supplementary Figure S8. A heat map showing the expression level of all *PLL* genes from PPGP II transcriptome data for *T. versicolor*, assembly BC4**. Note that *TrVePLL1* is missing in BC4, thus not included in this figure. Blue star on the right indicates this gene is differentially expressed in haustorial tissues. The yellow star on the right indicates the highest expression level of this gene is found in haustorial tissues.

## References

Abramson J, Adler J, Dunger J, Evans R, Green T, Pritzel A, Ronneberger O, Willmore L, Ballard AJ, Bambrick J, et al. 2024. Accurate structure prediction of biomolecular interactions with AlphaFold 3. Nature 630: 493–500.

Aly R, Cholakh H, Joel DM, Leibman D, Steinitz B, Zelcer A, Naglis A, Yarden O, Gal-On A. 2009. Gene silencing of mannose 6-phosphate reductase in the parasitic weed Orobanche aegyptiaca through the production of homologous dsRNA sequences in the host plant. Plant biotechnology journal 7: 487–498.

Auriac M-C, Griffiths C, Robin-Soriano A, Legendre A, Boniface M-C, Muños S, Fournier J, Chabaud M. 2024. The penetration of sunflower root tissues by the parasitic plant Orobanche cumana is intracellular. The new phytologist 241: 2326–2332.

Bandaranayake PCG, Filappova T, Tomilov A, Tomilova NB, Jamison-McClung D, Ngo Q, Inoue K, Yoder JI. 2010. A single-electron reducing quinone oxidoreductase is necessary to induce haustorium development in the root parasitic plant Triphysaria. The Plant Cell Online 22: 1404–1419.

Bandaranayake PC, Yoder JI. 2013. Trans-specific gene silencing of acetyl-CoA carboxylase in a root-parasitic plant. Molecular plant-microbe interactions: MPMI 26: 575–584.

Bandaranayake PCG, Yoder JI. 2018. Factors affecting the efficiency of Rhizobium rhizogenes root transformation of the root parasitic plant Triphysaria versicolor and its host Arabidopsis thaliana. Plant methods 14: 61.

Bawin T, Didriksen A, Faehn C, Olsen S, Sørensen I, Rose JKC, Krause K. 2023. Cuscuta campestris fine-tunes gene expression during haustoriogenesis as an adaptation to different hosts. Plant physiology 194: 258–273.

Bellis ES, Kelly EA, Lorts CM, Gao H, DeLeo VL, Rouhan G, Budden A, Bhaskara GB, Hu Z, Muscarella R, et al. 2020. Genomics of sorghum local adaptation to a parasitic plant. Proceedings of the National Academy of Sciences of the United States of America 117: 4243– 4251.

Boisson-Dernier A, Chabaud M, Garcia F, Bécard G, Rosenberg C, Barker DG. 2001. Agrobacterium rhizogenes-transformed roots of Medicago truncatula for the study of nitrogen-fixing and endomycorrhizal symbiotic associations. Molecular plant-microbe interactions: MPMI 14: 695–700.

Bradley JM, Butlin RK, Scholes JD. 2024. Comparative secretome analysis of Striga and Cuscuta species identifies candidate virulence factors for two evolutionarily independent parasitic plant lineages. BMC plant biology 24: 251.

Camacho C, Coulouris G, Avagyan V, Ma N, Papadopoulos J, Bealer K, Madden TL. 2009. BLAST+: architecture and applications. BMC bioinformatics 10: 421.

Capella-Gutiérrez S, Silla-Martínez JM, Gabaldón T. 2009. trimAl: a tool for automated alignment trimming in large-scale phylogenetic analyses. Bioinformatics 25: 1972–1973.

Collmer A, Keen NT. 1986. The role of pectic enzymes in plant pathogenesis. Annual Review of Phytopathology. 24:383–409

Crooks GE, Hon G, Chandonia J-M, Brenner SE. 2004. WebLogo: a sequence logo generator. Genome research 14: 1188–1190.

Cui S, Wakatake T, Hashimoto K, Saucet SB, Toyooka K, Yoshida S, Shirasu K. 2016. Haustorial Hairs Are Specialized Root Hairs That Support Parasitism in the Facultative Parasitic Plant Phtheirospermum japonicum. Plant physiology 170: 1492–1503.

Dashek WV, Rosen WG. 1966. Electron microscopical localization of chemical components in the growth zone of lily pollen tubes. Protoplasma 61: 192–204.

Dixit S, Upadhyay SK. 2023. Melting the wall: plant parasitism entails pectin modification. Plant signaling & behavior 18: 2252219.

Eddy SR. 2011. Accelerated Profile HMM Searches. PLoS computational biology 7: e1002195.

Eticha D, Stass A, Horst WJ. 2005. Cell-wall pectin and its degree of methylation in the maize root-apex: significance for genotypic differences in aluminium resistance. Plant, cell & environment 28: 1410–1420.

Fernández-Aparicio M, Delavault P, Timko MP. 2020. Management of Infection by Parasitic Weeds: A Review. Plants 9: 1184

de Framond A, Rich PJ, McMillan J, Ejeta G. 2007. Effects on Striga parasitism of transgenic maize armed with RNAi constructs targeting essential S. asiatica genes. In: Integrating New Technologies for Striga Control. WORLD SCIENTIFIC, 185–196.

Gaffe J, Tieman DM, Handa AK. 1994. Pectin Methylesterase Isoforms in Tomato (Lycopersicon esculentum) Tissues (Effects of Expression of a Pectin Methylesterase Antisense Gene). Plant physiology 105: 199–203.

Gao F, Chen C, Arab DA, D. Z, He Y, Ho SYW. 2019. EasyCodeML: A visual tool for analysis of selection using CodeML. Ecology and evolution 9: 3891–3898.

Gressel J, Gassmann AJ, Owen MD. 2017. How well will stacked transgenic pest/herbicide resistances delay pests from evolving resistance? Pest management science 73: 22–34.

Haas BJ. 2024. Transdecoder. GitHub.

Heslop-Harrison J, Heslop-Harrison Y. 1982. The growth of the grass pollen tube: 1. Characteristics of the polysaccharide particles (‘P-particles’) associated with apical growth. Protoplasma 112: 71–80.

Holt C, Yandell M. 2011. MAKER2: an annotation pipeline and genome-database management tool for second-generation genome projects. BMC bioinformatics 12: 491.

Hoshino A, Jayakumar V, Nitasaka E, Toyoda A, Noguchi H, Itoh T, Shin-I T, Minakuchi Y, Koda Y, Nagano AJ, et al. 2016. Genome sequence and analysis of the Japanese morning glory Ipomoea nil. Nature communications 7: 13295.

Ischebeck T, Stenzel I, Heilmann I. 2008. Type B phosphatidylinositol-4-phosphate 5-kinases mediate Arabidopsis and Nicotiana tabacum pollen tube growth by regulating apical pectin secretion. The Plant cell 20: 3312–3330.

Jhu M-Y, Farhi M, Wang L, Zumstein K, Sinha NR. 2021a. Investigating Host and Parasitic Plant Interaction by Tissue-Specific Gene Analyses on Tomato and Cuscuta campestris Interface at Three Haustorial Developmental Stages. Frontiers in plant science 12: 764843.

Jhu M-Y, Ichihashi Y, Farhi M, Wong C, Sinha NR. 2021b. LATERAL ORGAN BOUNDARIES DOMAIN 25 functions as a key regulator of haustorium development in dodders. Plant physiology 186: 2093–2110.

Jiang J, Yao L, Yu Y, Liang Y, Jiang J, Ye N, Miao Y, Cao J. 2014. PECTATE LYASE-LIKE 9 from Brassica campestris is associated with intine formation. Plant science: an international journal of experimental plant biology 229: 66–75.

John P. Vogel, Theodore K. Raab, Celine Schiff, Shauna C. Somerville. 2002. PMR6, a Pectate Lyase–Like Gene Required for Powdery Mildew Susceptibility in Arabidopsis. The Plant Cell 14: 2095–2106.

Jones P, Binns D, Chang H-Y, Fraser M, Li W, McAnulla C, McWilliam H, Maslen J, Mitchell A, Nuka G, et al. 2014. InterProScan 5: genome-scale protein function classification. Bioinformatics 30: 1236–1240.

Katoh K, Misawa K, Kuma K, Miyata T. 2002. MAFFT: a novel method for rapid multiple sequence alignment based on fast Fourier transform. Nucleic acids research 30: 3059–3066.

Koressaar T, Remm M. 2007. Enhancements and modifications of primer design program Primer3. Bioinformatics 23: 1289–1291.

Kotoujansky A. 1987. Molecular Genetics of Pathogenesis by Soft-Rot Erwinias. Annual review of phytopathology 25: 405–430.

Krogh A, Larsson B, von Heijne G, Sonnhammer EL. 2001. Predicting transmembrane protein topology with a hidden Markov model: application to complete genomes. Journal of molecular biology 305: 567–580.

Kurotani K-I, Wakatake T, Ichihashi Y, Okayasu K, Sawai Y, Ogawa S, Cui S, Suzuki T, Shirasu K, Notaguchi M. 2020. Host-parasite tissue adhesion by a secreted type of β-1,4-glucanase in the parasitic plant Phtheirospermum japonicum. Communications biology 3: 407.

Leso M, Kokla A, Feng M, Melnyk CW. 2023. Pectin modifications promote haustoria development in the parasitic plant Phtheirospermum japonicum. Plant physiology 194: 229– 242.

Livak KJ, Schmittgen TD. 2001. Analysis of relative gene expression data using real-time quantitative PCR and the 2(-Delta Delta C(T)) Method. Methods (San Diego, Calif.) 25: 402–408.

Li Y, Wei W, Feng J, Luo H, Pi M, Liu Z, Kang C. 2018. Genome re-annotation of the wild strawberry Fragaria vesca using extensive Illumina-and SMRT-based RNA-seq datasets. DNA research: an international journal for rapid publication of reports on genes and genomes 25: 61– 70.

Losner-Goshen D, Portnoy VH, Mayer AM, Joel DM. 1998. Pectolytic Activity by the Haustorium of the Parasitic PlantOrobancheL. (Orobanchaceae) in Host Roots. Annals of botany 81: 319–326.

Marx H, Minogue CE, Jayaraman D, Richards AL, Kwiecien NW, Siahpirani AF, Rajasekar S, Maeda J, Garcia K, Del Valle-Echevarria AR, et al. 2016. A proteomic atlas of the legume Medicago truncatula and its nitrogen-fixing endosymbiont Sinorhizobium meliloti. Nature biotechnology 34: 1198–1205.

McCarthy TW, Der JP, Honaas LA, dePamphilis CW, Anderson CT. 2014. Phylogenetic analysis of pectin-related gene families in Physcomitrella patensand nine other plant species yields evolutionary insights into cell walls. BMC plant biology 14: 1–14.

Minh BQ, Nguyen MAT, von Haeseler A. 2013. Ultrafast approximation for phylogenetic bootstrap. Molecular biology and evolution 30: 1188–1195.

Minh BQ, Schmidt HA, Chernomor O, Schrempf D, Woodhams MD, von Haeseler A, Lanfear R. 2020. IQ-TREE 2: New Models and Efficient Methods for Phylogenetic Inference in the Genomic Era. Molecular biology and evolution 37: 1530–1534.

Nakajima N, Ishihara K, Tanabe M, Matsubara K, Matsuura Y. 1999. Degradation of pectic substances by two pectate lyases from a human intestinal bacterium, Clostridium butyricum-beijerinckii group. Journal of bioscience and bioengineering 88: 331–333.

Ohtsuki T, Taniguchi Y, Kohno K, Fukuda S, Usui M, Kurimoto M. 1995. Cry j 2, a major allergen of Japanese cedar pollen, shows polymethylgalacturonase activity. Allergy 50: 483– 488.

Palusa S, Golovkin M, Shin S, Richardson DN, Reddy ASN. 2007. Organ-specific, developmental, hormonal and stress regulation of expression of putative pectate lyase genes in Arabidopsis. The New phytologist 174: 537–550.

Pilkington SM, Crowhurst R, Hilario E, Nardozza S, Fraser L, Peng Y, Gunaseelan K, Simpson R, Tahir J, Deroles SC, et al. 2018. A manually annotated Actinidia chinensis var. chinensis (kiwifruit) genome highlights the challenges associated with draft genomes and gene prediction in plants. BMC genomics 19: 257.

Porter RH, Durrell M, Romm HJ. 1947. The Use of 2,3,5-Triphenyl-Tetrazoliumchloride as a Measure of Seed Germinability. Plant physiology 22: 149–159.

Pradhan Mitra P, Loqué D. 2014. Histochemical staining of Arabidopsis thaliana secondary cell wall elements. Journal of visualized experiments: JoVE.

Ridley BL, O’Neill MA, Mohnen D. 2001. Pectins: structure, biosynthesis, and oligogalacturonide-related signaling. Phytochemistry 57: 929–967.

Rosen WG, Gawlik SR. 1966. Fine structure of lily pollen tubes following various fixation and staining procedures. Protoplasma 61: 181–191.

Runo S, Alakonya A, Machuka J, Sinha N. 2011. RNA interference as a resistance mechanism against crop parasites in Africa: a ‘Trojan horse’ approach. Pest management science 67: 129– 136.

Scavetta RD, Herron SR, Hotchkiss AT, Kita N, Keen NT, Benen JAE, Kester HCM, Visser J, Jurnak F. 1999. Structure of a Plant Cell Wall Fragment Complexed to Pectate Lyase C. The Plant cell 11: 1081–1092.

Schneider TD, Stephens RM. 1990. Sequence logos: a new way to display consensus sequences. Nucleic acids research 18: 6097–6100.

Schrödinger, LLC. 2015. The PyMOL Molecular Graphics System, Version 3.1.6.1

Stamatakis A. 2014. RAxML version 8: a tool for phylogenetic analysis and post-analysis of large phylogenies. Bioinformatics 30: 1312–1313.

Stanke M, Steinkamp R, Waack S, Morgenstern B. 2004. AUGUSTUS: a web server for gene finding in eukaryotes. Nucleic acids research 32: W309–12.

Starr MP, Moran F. 1962. Eliminative split of pectic substances by phytopathogenic soft-rot bacteria. Science 135: 920–921.

Steele DB. 2021. The Transcriptional Regulation of Host Recognition and Prehaustorium Development in Triphysaria versicolor. PhD dissertation, University of California, Davis.

Stūrīte I, Henriksen TM, Breland TA. 2005. Distinguishing between metabolically active and inactive roots by combined staining with 2,3,5-triphenyltetrazolium chloride and image colour analysis. Plant and soil 271: 75–82.

Sun H, Hao P, Gu L, Cheng S, Wang H, Wu A, Ma L, Wei H, Yu S. 2020. Pectate lyase-like Gene GhPEL76 regulates organ elongation in Arabidopsis and fiber elongation in cotton. Plant science: an international journal of experimental plant biology 293: 110395.

Sun H, Hao P, Ma Q, Zhang M, Qin Y, Wei H, Su J, Wang H, Gu L, Wang N, et al. 2018. Genome-wide identification and expression analyses of the pectate lyase (PEL) gene family in cotton (Gossypium hirsutum L.). BMC genomics 19: 661.

Sun L, van Nocker S. 2010. Analysis of promoter activity of members of the PECTATE LYASE-LIKE (PLL) gene family in cell separation in Arabidopsis. BMC plant biology 10: 152.

Teufel F, Almagro Armenteros JJ, Johansen AR, Gíslason MH, Pihl SI, Tsirigos KD, Winther O, Brunak S, von Heijne G, Nielsen H. 2022. SignalP 6.0 predicts all five types of signal peptides using protein language models. Nature biotechnology 40: 1023–1025.

The Galaxy Community. 2022. The Galaxy platform for accessible, reproducible and collaborative biomedical analyses: 2022 update. Nucleic acids research 50: W345–W351.

Tomilov A, Tomilova N, Yoder JI. 2006. Agrobacterium tumefaciens and Agrobacterium rhizogenes transformed roots of the parasitic plant Triphysaria versicolor retain parasitic competence. Planta 225: 1059–1071.

Uluisik S, Seymour GB. 2020. Pectate lyases: Their role in plants and importance in fruit ripening. Food chemistry 309: 125559.

Untergasser A, Cutcutache I, Koressaar T, Ye J, Faircloth BC, Remm M, Rozen SG. 2012. Primer3—new capabilities and interfaces. Nucleic acids research 40: e115–e115.

Vierheilig H, Coughlan AP, Wyss U, Piche Y. 1998. Ink and vinegar, a simple staining technique for arbuscular-mycorrhizal fungi. Applied and environmental microbiology 64: 5004– 5007.

Wafula EK, Zhang H, Von Kuster G, Leebens-Mack JH, Honaas LA, dePamphilis CW. 2022. PlantTribes2: Tools for comparative gene family analysis in plant genomics. Frontiers in plant science 13: 1011199.

Wang Y, Murdock M, Lai SWT, Steele DB, Yoder JI. 2020. Kin recognition in the parasitic plant Triphysaria versicolor is mediated through root exudates. Frontiers in plant science 11: 560682.

Weisenfeld NI, Kumar V, Shah P, Church DM, Jaffe DB. 2017. Direct determination of diploid genome sequences. Genome research 27: 757–767.

Westwood JH, dePamphilis CW, Das M, Fernández-Aparicio M, Honaas LA, Timko MP, Wafula EK, Wickett NJ, Yoder JI. 2012. The Parasitic Plant Genome Project: New Tools for Understanding the Biology of Orobanche and Striga. Weed Science 60: 295306.

Wu S, Lau KH, Cao Q, Hamilton JP, Sun H, Zhou C, Eserman L, Gemenet DC, Olukolu BA, Wang H, et al. 2018. Genome sequences of two diploid wild relatives of cultivated sweet potato reveal targets for genetic improvement. Nature communications 9: 4580.

Wu Y, Qiu X, Du S, Erickson L. 1996. PO149, a new member of pollen pectate lyase-like gene family from alfalfa. Plant molecular biology 32: 1037–1042.

Xu Z, Dai J, Kang T, Shah K, Li Q, Liu K, Xing L, Ma J, Zhang D, Zhao C. 2022. PpePL1 and PpePL15 Are the Core Members of the Pectate Lyase Gene Family Involved in Peach Fruit Ripening and Softening. Frontiers in plant science 13: 844055.

Yang Z. 2007. PAML 4: phylogenetic analysis by maximum likelihood. Molecular biology and evolution 24: 1586–1591.

Yang L, Huang W, Xiong F, Xian Z, Su D, Ren M, Li Z. 2017. Silencing of SlPL, which encodes a pectate lyase in tomato, confers enhanced fruit firmness, prolonged shelf-life and reduced susceptibility to grey mould. Plant biotechnology journal 15: 1544–1555.

Yang Z, Wafula EK, Honaas LA, Zhang H, Das M, Fernandez-Aparicio M, Huang K, Bandaranayake PC, Wu B, Der JP, et al. 2015. Comparative transcriptome analyses reveal core parasitism genes and suggest gene duplication and repurposing as sources of structural novelty. Molecular biology and evolution 32: 767–790.

Yang Z, Wong WSW, Nielsen R. 2005. Bayes empirical bayes inference of amino acid sites under positive selection. Molecular biology and evolution 22: 1107–1118.

Yocca A, Akinyuwa M, Bailey N, Cliver B, Estes H, Guillemette A, Hasannin O, Hutchison J, Jenkins W, Kaur I, et al. 2024. A chromosome-scale assembly for ‘d’Anjou’ pear. G3 (Bethesda, Md.) 14.

Yoder MD, Jurnak F. 1995. The Refined Three-Dimensional Structure of Pectate Lyase C from Erwinia chrysanthemi at 2.2 Angstrom Resolution (Implications for an Enzymatic Mechanism). Plant physiology 107: 349–364.

Yoshida S, Cui S, Ichihashi Y, Shirasu K. 2016. The Haustorium, a Specialized Invasive Organ in Parasitic Plants. Annual review of plant biology 67: 643–667.

Yujia Leng, Yaolong Yang, Deyong Ren, Lichao Huang, Liping Dai, Yuqiong Wang, Long Chen, Zhengjun Tu, Yihong Gao, Xueyong Li, Li Zhu, Jiang Hu, Guangheng Zhang, Zhenyu Gao, Longbiao Guo, Zhaosheng Kong, Yongjun Lin, Qian Qian, Dali Zeng. 2017. A rice PECTATE LYASE-LIKE gene is required for plant growth and leaf senescence. Plant Physiology 174: 1151–1166.

Zhang H. 2020. Phylogenetic and functional studies on parasitism genes in Orobanchaceae and Convolvulaceae (C dePamphilis, Ed.).

Zhang H, Wafula EK, Eilers J, Harkess AE, Ralph PE, Timilsena PR, dePamphilis CW, Waite JM, Honaas LA. 2022. Building a foundation for gene family analysis in Rosaceae genomes with a novel workflow: A case study in Pyrus architecture genes. Frontiers in plant science 13: 975942.

Zheng Y, Yan J, Wang S, Xu M, Huang K, Chen G, Ding Y. 2018. Genome-wide identification of the pectate lyase-like (PLL) gene family and functional analysis of two PLL genes in rice. Molecular genetics and genomics: MGG 293: 1317–1331.

