## supplemental figures for "The evolution of pectate lyase-like genes across land plants, and their roles in haustorium formation in parasitic plant, *Triphysaria versicolor* (Orobanchaceae)"

Supplemental Figure S1.

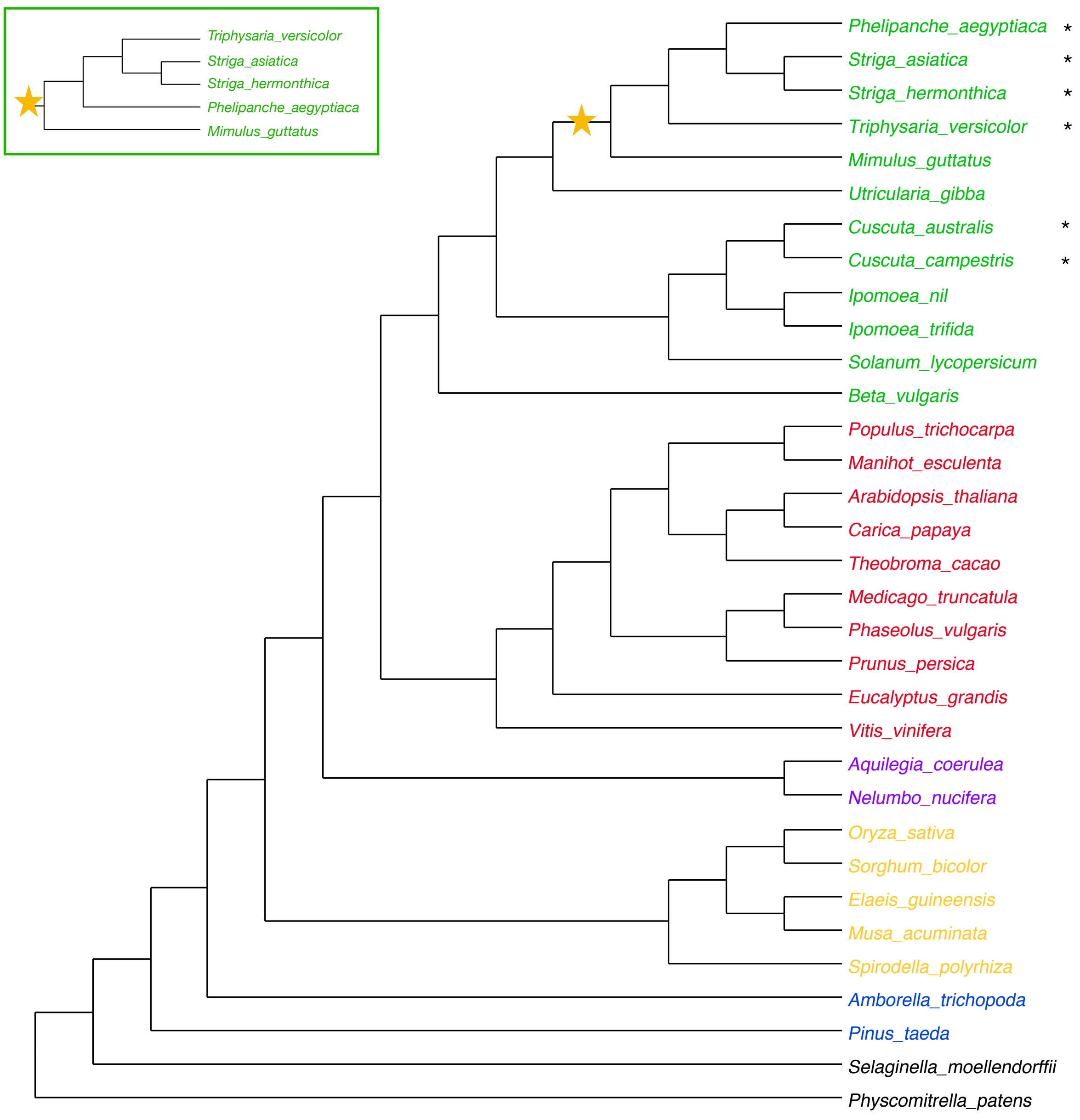

Supplemental Figure S2.

A

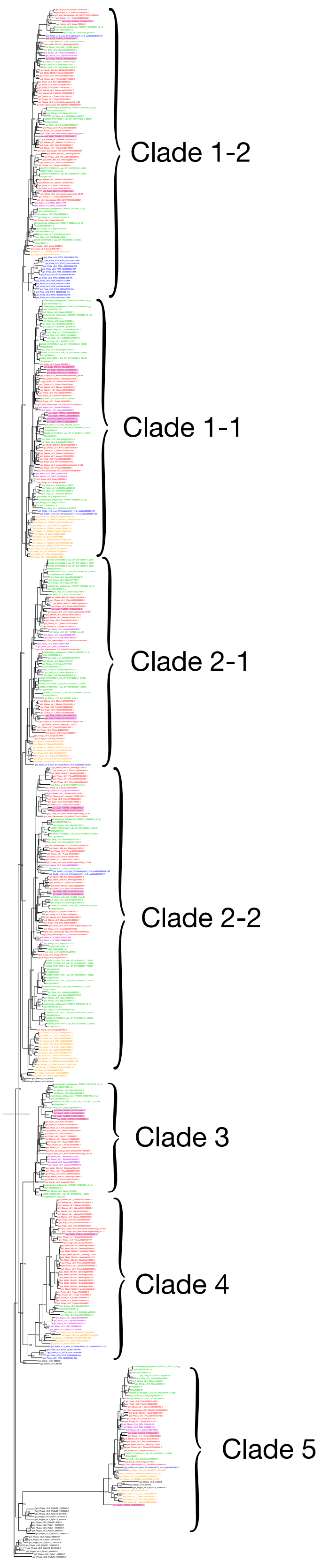

B

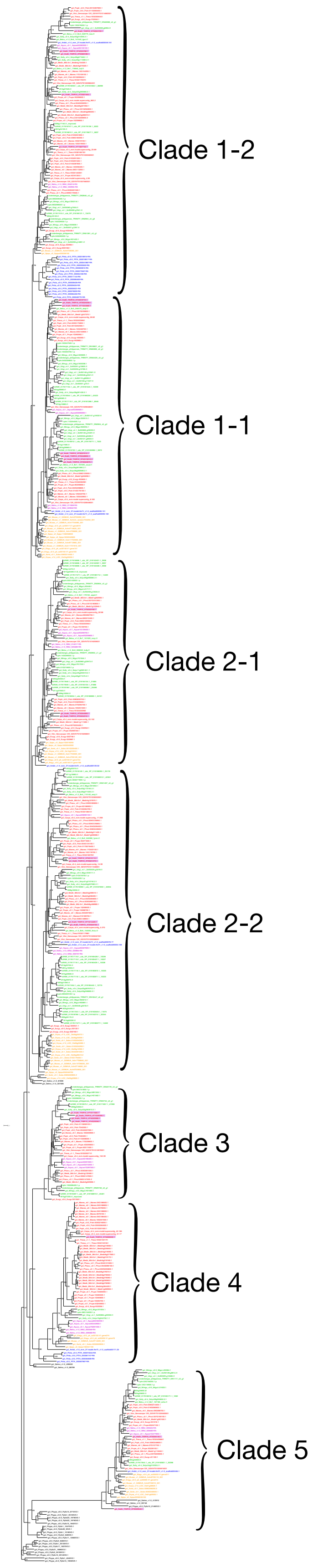

C

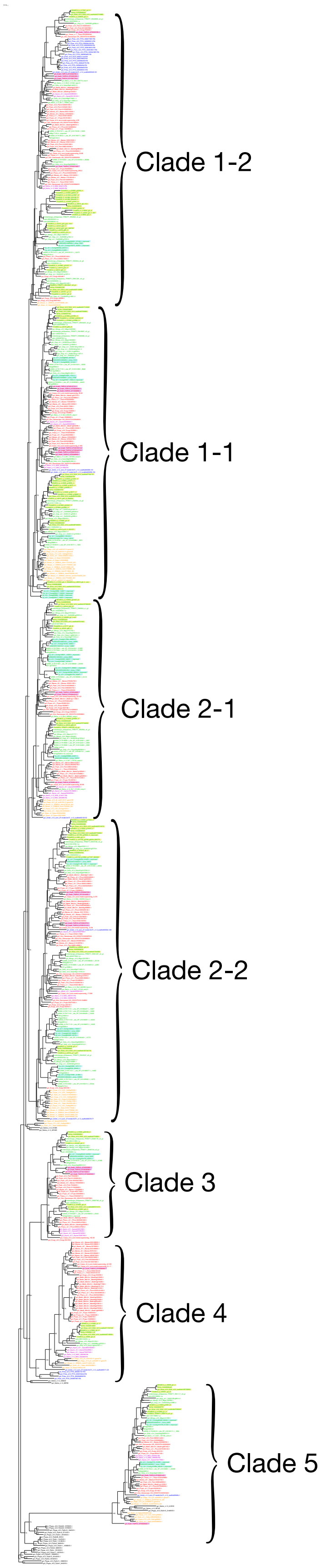

D

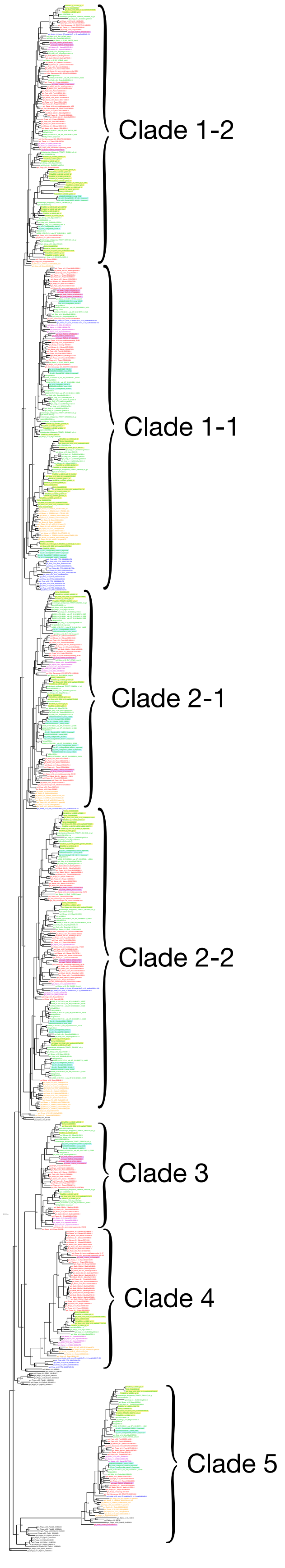

Supplemental Figure S3.

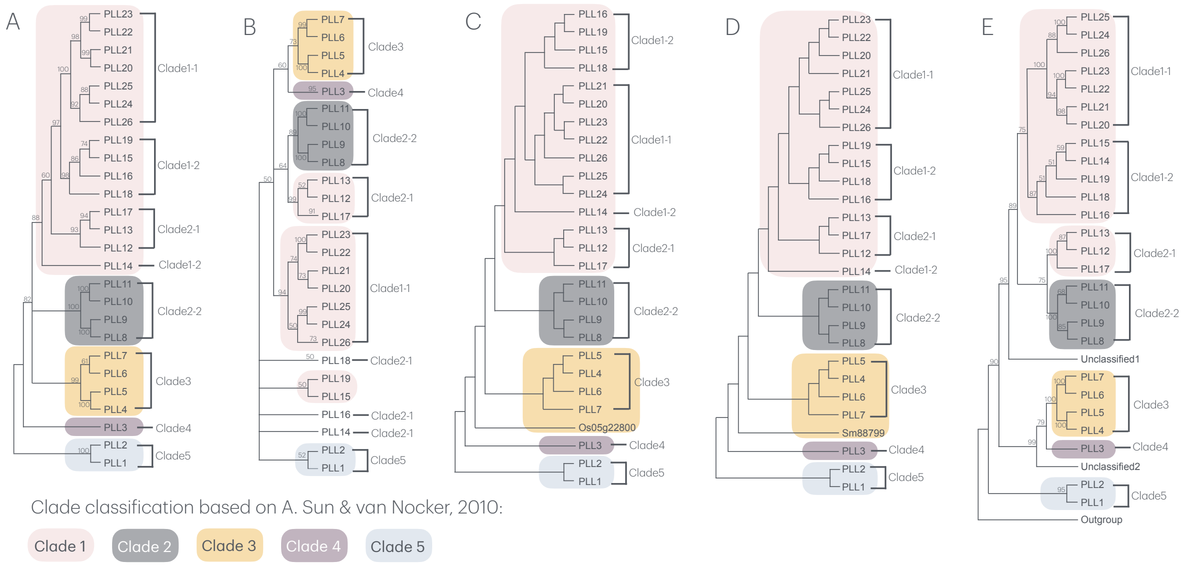


| F | Phylogeny A | Phylogeny B | Phylogeny C | Phylogeny D | Phylogeny from this study E |
|---|---|---|---|---|---|
| **Source** | Sun and van Nocker, 2010 | McCarthy et al., 2014 | Sun et al., 2018 | Zheng et al., 2018 | This study |
| **Sampling strategy** | Arabidopsis PLL genes only, identified via a BLAST search querying a banana PL gene | PLLs identified via a orthogroup approach, querying Arabidopsis PLLs in 10 diverse land plant genomes | PLLs identified from 7 diverse land plants (including 3 grasses) via BLAST searches | PLLs identified from 3 *Gossypium* species via HMMER search querying the pec_lyase_C domain. Method identifying PLLs from other plant species is missing. | PLLs identified via a orthogroup approach, querying Arabidopsis PLLs in 36 diverse land plant genomes or transcriptomes. |
| **Alignment method** | Protein alignment with ClustalW2 | Protein alignment with MUSCLE | Protein alignment with ClustalX | Protein alignment with ClustalX 2.0 | Codon alignment with MAFFT |
| **Phylogeny inference method** | Neighbor-joining (MEGA4) | Maximum likelihood (RAxML) | Neighbor-joining (MEGA6.0) | Neighbor-joining (MEGA6.0) | Maximum likelihood (RAxML& IQtree) |
| **Bootstrap replicates** | Unknown | 1000 | 1000 | 1000 | 1000 for RAxML; 2000 for IQtree |
| **Notes** | • First attempt to study the PLL family in plants.<br>• Only one species is used.<br>• Method is not robust. | • Diverse selection of plant genomes.<br>• Robust methods.<br>• Multiple unresolved (bs<50) clades | • Increased but limited selection of plant genomes.<br>• Paraphyletic group recognized as a clade.<br>• No bootstrap shown<br>• Method is not robust | • Increased but limited selection of plant genomes.<br>• Paraphyletic group recognized as a clade.<br>• No bootstrap shown<br>• Method is not robust | • Diverse selection of plant genomes.<br>• Robust methods.<br>• Well resolved clades |

Supplemental Figure S4.

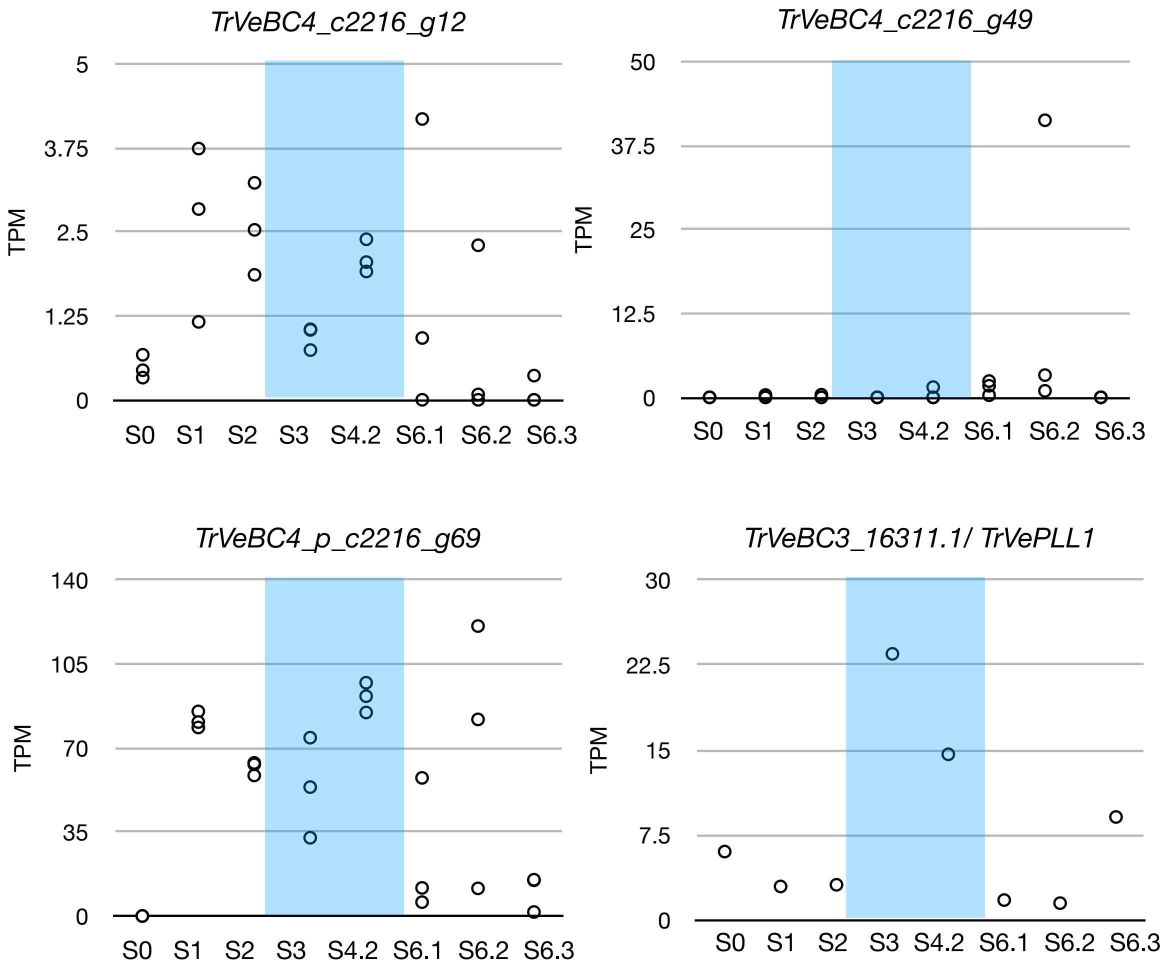

Supplemental Figure S5.

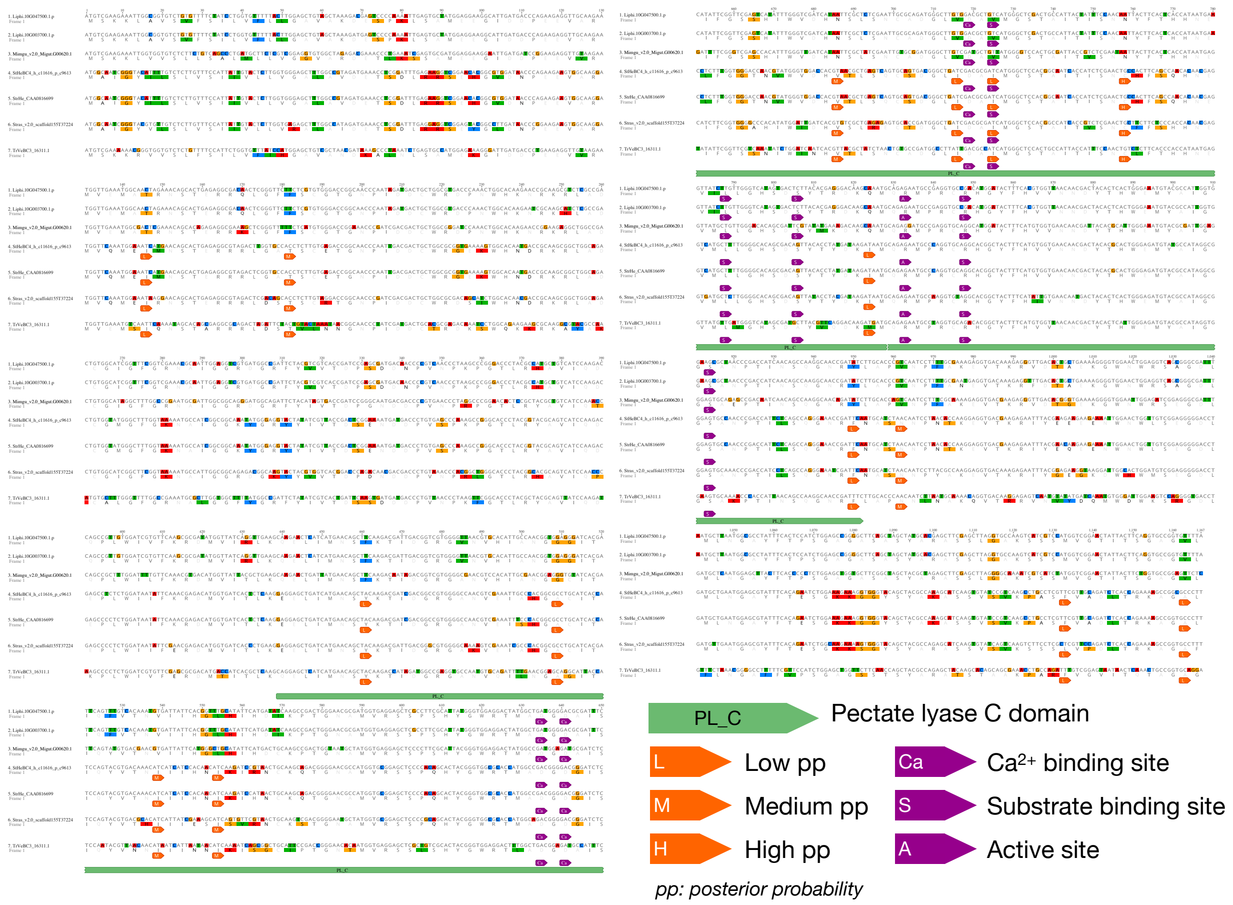

Supplemental Figure S6.

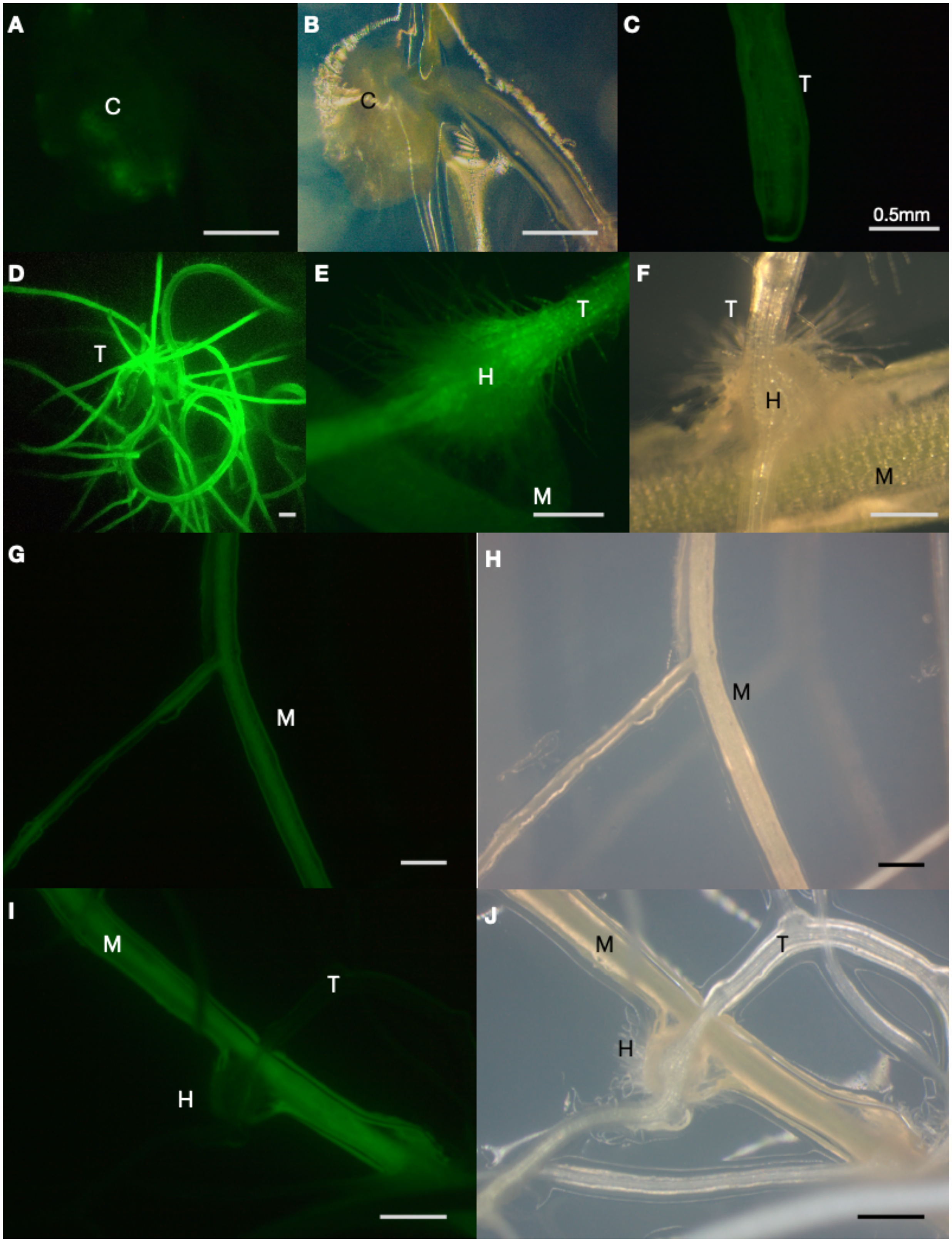

Supplemental Figure S7.

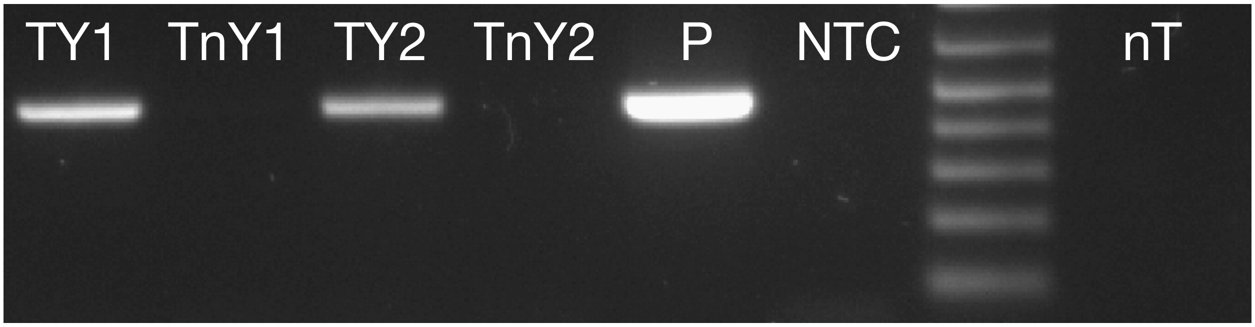

Supplemental Figure S8.

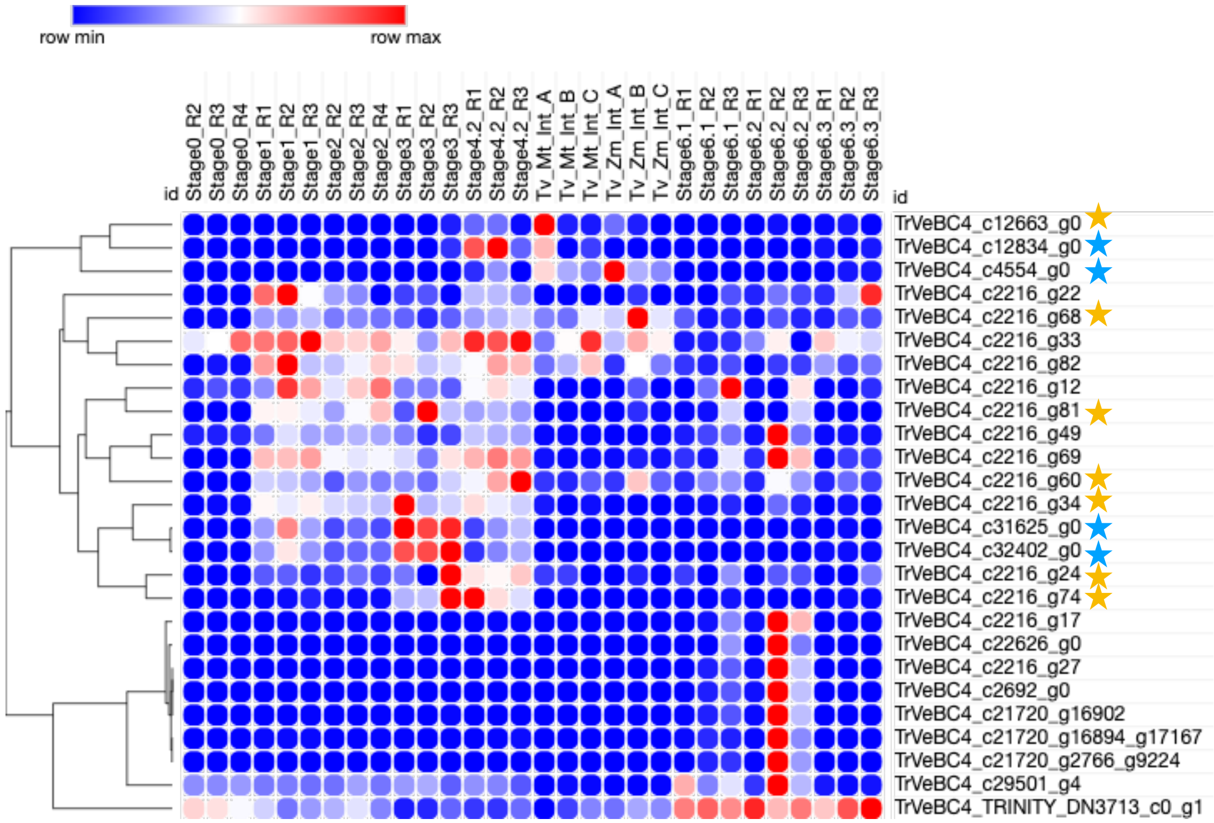
